# Structure and autoregulation of a P4-ATPase lipid flippase

**DOI:** 10.1101/606061

**Authors:** Milena Timcenko, Joseph A. Lyons, Dovile Januliene, Jakob Ulstrup, Thibaud Dieudonne, Cedric Montigny, Miriam Rose Ash, Jesper Lykkegaard Karlsen, Thomas Boesen, Werner Kühlbrandt, Guillaume Lenoir, Arne Möller, Poul Nissen

**Affiliations:** DANDRITE - Nordic EMBL Partnership for Molecular Medicine, Aarhus University, Dept. Molecular Biology and Genetics, Denmark; Max Planck Institute for Biophysics, Frankfurt; Institute for Integrative Biology of the Cell (I2BC), CEA, CNRS, Univ. Paris-Sud, Université Paris-Saclay, F-91198, Gif-sur-Yvette, France; Interdisciplinary Nanoscience Center - iNANO, Aarhus University

## Abstract

P4-ATPases are lipid flippases that drive active transport of phospholipids from the exoplasmic or lumenal to the cytosolic leaflets of eukaryotic membranes to maintain their asymmetric lipid composition. The molecular architecture of P4-ATPases and how they work in lipid recognition and transport has remained elusive. Using cryo-electron microscopy we have determined the structures of a P4-ATPase, specifically of the *Saccharomyces cerevisiae* Drs2p-Cdc50p, which is a phosphatidylserine and phosphatidylethanolamine specific lipid flippase. Drs2p-Cdc50p is autoinhibited by the Drs2p C-terminal tail and activated by phosphatidylinositol-4 phosphate (PI4P). We present three structures representing an autoinhibited, an intermediate, and a fully activated state. The analysis highlights specific features of P4-ATPases and reveals sites of auto-inhibition and PI4P-dependent activation. We observe the opening of a putative flippase pathway engaging conserved residues Ile508 of transmembrane segment 4 and Lys1018 and polar residues of transmembrane segment 5 in the centre of the lipid bilayer.

## Introduction

Cells and organelles are defined by lipid bilayer membranes and membrane proteins. Eukaryotic membranes of the secretory/endocytic pathways are typically asymmetric with respect to lipid distribution in the inner and outer bilayer leaflets. The resulting gradients in lipid concentration potentiate important biological processes such as membrane dynamics in morphological changes and motility of the cell, endo- and exocytosis, and signalling^1–4^. Due to membrane-fusion events and the activity of lipid scramblases, which move lipids between the inner and outer leaflet in both directions, lipid asymmetry must constantly be regulated and maintained. Members of two distinct membrane protein superfamilies drive the ATP-dependent unidirectional translocation of lipids against concentration gradients in the membrane; namely ATP-binding cassette (ABC) transporters and P-type ATPases of P4 subtype that generally drive an inward-to-outward (flop) and outward-to-inward (flip) translocation of lipids between bilayer leaflets, respectively^1–4^. P-type ATPases couple transport to ATP hydrolysis via formation and breakdown of a phosphoenzyme in a functional cycle with so-called E1, E1P, E2P and E2 intermediate states. The P4-ATPases specifically couple their lipid flippase activity to dephosphorylation of the E2P state^5,6^, i.e. similar to the inward K^+^ transport of Na,K-ATPase, whereas E1P phosphoenzyme formation seems independent of substrate binding^6,7^. While recent studies have shed light on the structure and function of lipid floppases^8,9^ and scramblases^10,11^, P4-ATPases have so far been studied only by bioinformatics and functional assays. To date, the transport mechanism remains enigmatic and is much debated, with models implicating either peripheral or centrally located lipid recognition sites and pathways^12–14^.

Most P4-ATPases are binary complexes, where a Cdc50-protein subunit is necessary for proper localization of the complex and probably also for function^15,16^. Mutant forms of mammalian lipid flippases have been implicated in disease, e.g. ATP8A1 and ATP8A2 in neurological disorders, ATP8B1 in progressive familial intrahepatic cholestasis type 1 (PFIC1), ATP10A in type 2 diabetes and insulin resistance, and ATP11A in cancer^17^. One of the best-characterized P4-ATPases is the trans-Golgi localized Drs2p-Cdc50p complex from the yeast *Saccharomyces cerevisiae*. *In vivo*^18,19^ and *in vitro*^20,21^ studies show that Drs2p-Cdc50p primarily flips phosphatidylserine (PS) and to a lesser extent phosphatidylethanolamine (PE) from the lumenal to the cytosolic leaflet, and indicate a role of this function in vesicle biogenesis at late secretory membranes.

The C-terminus of Drs2p contains an auto-inhibitory domain^22,23^. Relief of auto-inhibition requires the regulatory, but non-substrate lipid phosphatidylinositol 4-phosphate (PI4P)^5,22^. Binding of Gea2p (a guanine nucleotide exchange factor for the small GTPase Arf) to a basic segment of the C-terminus has been reported to be necessary for activation *in vivo*^22^, although this is not supported by studies *in vitro*^23^. Furthermore, interaction of Arl1p (another GTPase of the Arf family) with the extended N-terminus of Drs2p has been implicated in flippase activity *in vivo*^24^. Whereas the first 104 amino acids of the N-terminus have little effect on *in vitro* activity, truncation of the C-terminus has a strong activating effect, but the protein remains under regulation by PI4P^25^. While these studies highlight the components involved, the detailed molecular mechanism of autoregulation for Drs2p-Cdc50p and for P-type ATPases, in general, remains unknown.

To determine the structure of a P4-ATPase lipid flippase and investigate the molecular mechanism of transport and autoregulation, we embarked on cryo-EM studies of the beryllium fluoride (BeF_3_^−^) stabilized Drs2p-Cdc50p complex. Drs2p-Cdc50p was over-expressed in *S. cerevisiae* and purified in the detergent LMNG by affinity chromatography and gel filtration, resulting in a monodisperse sample containing both subunits (Supplementary Data Figure 1). Samples represent E2P phosphoenzyme-like states in the progressive steps from a fully autoinhibited P4-ATPase (E2P^inhib^), to an intermediate activated state in the presence of PI4P (E2P^inter^), and an outward-open and activated conformation captured using a C-terminally truncated enzyme also in the presence of PI4P (E2P^active^). The structures reveal the molecular architecture, pinpoint functional sites, and elucidate the PI4P-dependent regulation of Drs2p-Cdc50p.

### Overall structure and conformation

The structures of Drs2p-Cdc50p in the E2P^inhib^, E2P^inter^, and E2P^active^ conformations were determined at 2.8, 3.7 and 2.9 Å resolution, respectively (Supplementary Data Table 1). The density maps allowed complete modelling of the complexes with only some minor disordered regions at the termini missing.

The structure of the Drs2p subunit is typical of P-type ATPases with a transmembrane domain consisting of 10 helices, and three cytosolic domains: the actuator (A) domain, the nucleotide-binding (N) domain and the phosphorylation (P) domain (Figure 1A and B, Supplementary Figure 2A). The Cdc50p subunit has an ectodomain that folds into two asymmetric lobes where the first is dominated by an antiparallel β-sandwich and the other contains little secondary structure apart from short helical segments (Supplementary Data Figure 3A). Four glycosylation sites are apparent in the maps, with at least one *N*-acetylglucosamine unit resolved for each site in the final model. Two di-sulfide bonds identified earlier^26^ are evident (Supplementary Data Figure 3B). The fold of the ecto-domain is similar to the lipid binding protein seipin, although loops of Cdc50p are considerably longer (Supplementary Data Figure 3C). The two TM-helices of Cdc50p extend from the N- and C-terminus of the first lobe (Supplementary Data Figure 3A), interacting closely with each other and with TM10 of Drs2p (Supplementary Data Figure 3D).

**Figure 1:**
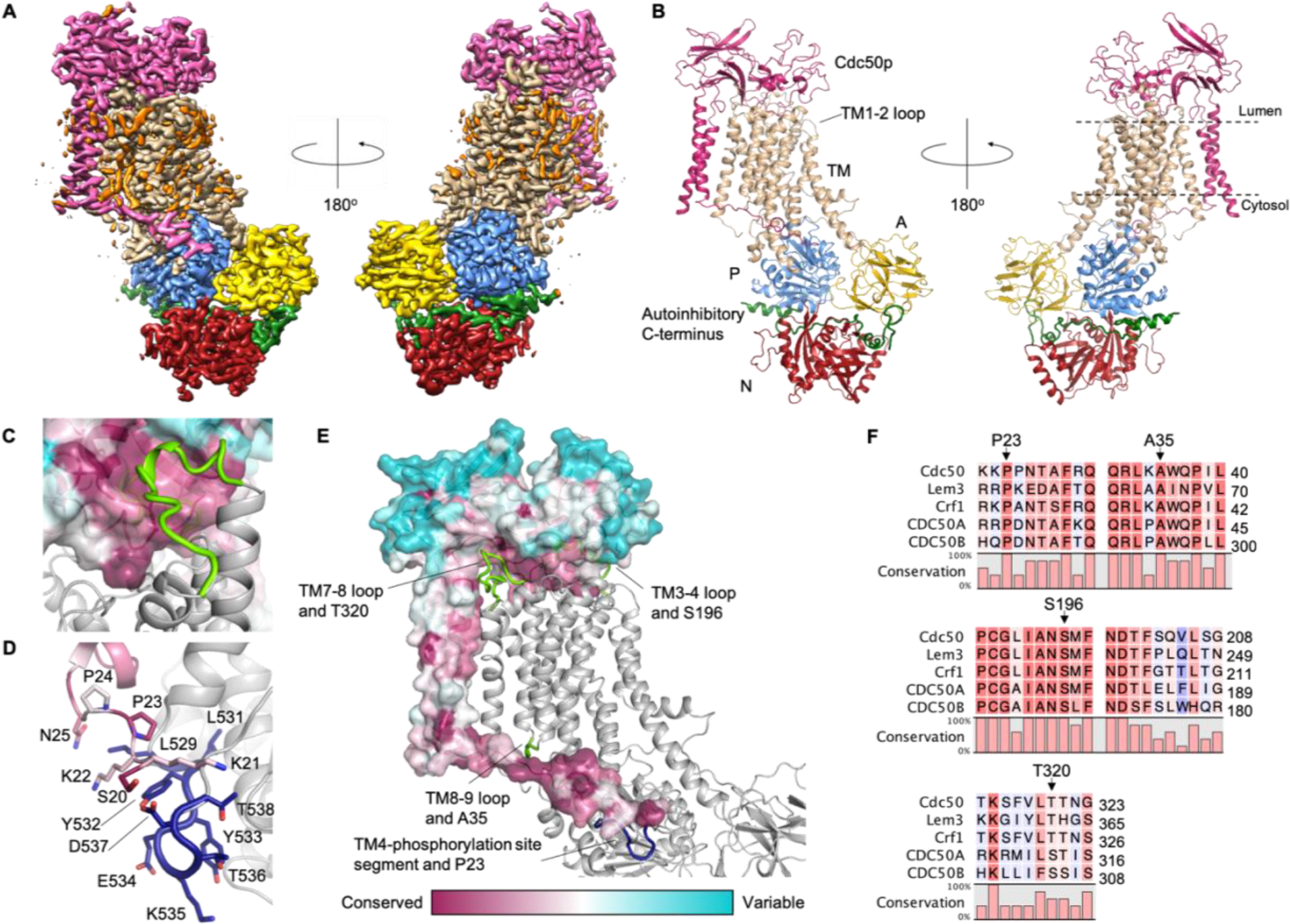
The structure of the Drs2p-Cdc50p complex. A) LocScale map^37^ of E2P^inhib^ colored by domain. For Drs2p the transmembrane (TM) domain is tan, the actuator (A) domain is yellow, the phosphorylation (P) domain is blue, the nucleotide binding (N) domain is red, and the autoinhibitory C-terminus is green. Cdc50p is pink. Unmodeled map features corresponding to ordered detergent molecules or from the detergent micelle are orange. B) Structural cartoon of the refined E2P^inhib^ model, c.f.A) C) Interaction between the Cdc50p ecto-domain (shown as surface) and the lumenal TM3-4 loop of Drs2p (light green). D)Interaction between the N-terminus of Cdc50p and the segment of Drs2p leading from TM4 to the phosphorylation site. Residues 529-538 are not present in P2 ATPases and are shown in dark blue. E)Segments of Drs2p found to interact with Cdc50p mutants that disrupt complex formation are highlighted in green and the insert in Drs2p between TM4 and the phosphorylation site is blue. The structure shown in C-E is E2P^inhib^, and Cdc50p is colored by conservation using ConSurf^38^. F)Part of a sequence alignment of Cdc50-proteins from *S. cerevisiae* and human CDC50A and CDC50B with residues important for complex-formation identified. Uniprot identifiers: Cdc50 – P25656, Lem3 – P42838, Crf1 – P53740, CDC50A – Q9NV96, CDC50B – Q3MIR4. The alignment was performed using Clustal Omega^39,40^.

#### Interactions between Drs2p and Cdc50p

The three structures show an invariant, tight complex between Drs2p and Cdc50p with interactions on the cytosolic and lumenal side and in the membrane. The most extensive interactions appear on the lumenal face of Drs2p, where the ecto-domain of Cdc50p interacts with all lumenal loops apart from the TM1-2 loop (Figure 1B). The TM3-4 loop in particular stretches into an intimate interaction site at the ectodomain of Cdc50p (Figure 1C). Chimeras between Drs2p and Dnf1p of this loop result in an intact but non-functional flippase^27^. Contacts in the transmembrane regions of the two proteins appear to be fewer, as only TM10 of Drs2p interacts with the helices of Cdc50p (Supplementary Data Figure 3D). On the cytosolic side, the N-terminus of Cdc50p wraps around the TM-domain of Drs2p and makes contacts with the segment (residues 529-538) that connects TM4 and the phosphorylation site of the P-domain (Figure 1D). This segment is 10 residues longer in P4-ATPases than in the P2-ATPase ion pumps, where TM4 couples the chemistry at the phosphorylation site with conformational changes of the TM-domain^28^. Interestingly, the TM4 segment as well as the TM3-4 loop are distinctly shorter in Neo1p and homologous lipid flippases that do not bind Cdc50p-subunits (Supplementary Data Figure 4). These interactions suggest that Cdc50p is sensitive to the phosphorylation state of the P-domain on the cytosolic side, while on the lumenal side it will sense the conformation of the transmembrane domain through the contact with the TM3-4 loop and other lumenal loops.

Mutations disrupting the interaction between Drs2p and Cdc50p have been identified in both the N-terminus of Cdc50p and in the interface with the TM3-4 loop of Drs2p (Figure 1E-F). In the *S. cerevisiae* phosphatidylcholine (PC) transporting Dnf1p-Lem3p, a Ser237Leu mutation in Lem3p (Ser196 in Cdc50p, located at the interface with the TM3-4 loop of Drs2p) disrupted the interaction between the two subunits, and Ala65Val (corresponding to Ala35 in Cdc50p near the TM8-9 loop of Drs2p), resulted in a mild reduction of interaction between the two subunits^26^. Furthermore, combining Pro23His and Thr320Ala mutations in Cdc50p resulted rendered growth in a *Δcdc50Δlem3Δcrf1* background temperature-sensitive^29^. These mutations are located at the Cdc50p N-terminus approaching the cytoplasmic end of Drs2p-TM4 and within the ecto-domain of Cdc50p (Figure 1E).

#### Comparison to other P-type ATPases

Drs2p-Cdc50p can be compared to Na,K-ATPase, which is also a P-type ATPase forming an αβ binary complex. However, the fold of Cdc50p and its discrete interactions in the membrane and with the lumenal face of Drs2p are fundamentally different from the β-subunit and regulatory FXYD subunit of Na,K-ATPase. The overall conformation of Drs2p is close to the ouabain-inhibited E2P-form of Na,K-ATPase (PDB 4HYT^30^) and the outward-open E2-BeF_3_^−^ form of SERCA (PDB 3B9B^31^). E2P^active^ resembles 3B9B most closely, with an overall root mean square deviation (rmsd) of 4.4 Å (superposition of C_α_-atoms, excluding the N-domain, which is flexible) compared to the E2P^inter^ and E2P^inhib^ conformations with a rmsd of 4.8 Å and 5.2 Å, respectively (Figure 2A). Other P-type ATPase conformations differ by a further margin of at least 0.8 Å in rmsd. The bulk of the A-domain is shifted by about 7 Å, but the (T/D)GE(S/T) loops of SERCA and Drs2p overlap, and density for the phosphate analogue BeF3 and bound Mg^2+^ is readily observable (Figure 2B). The conformation of the dephosphorylation loops are similar (Figure 2C). Compared to the ion-transporting P-type ATPases, TM1 and 2 appear to be one turn shorter at the lumenal side, suggesting they sit deeper in the membrane.

**Figure 2:**
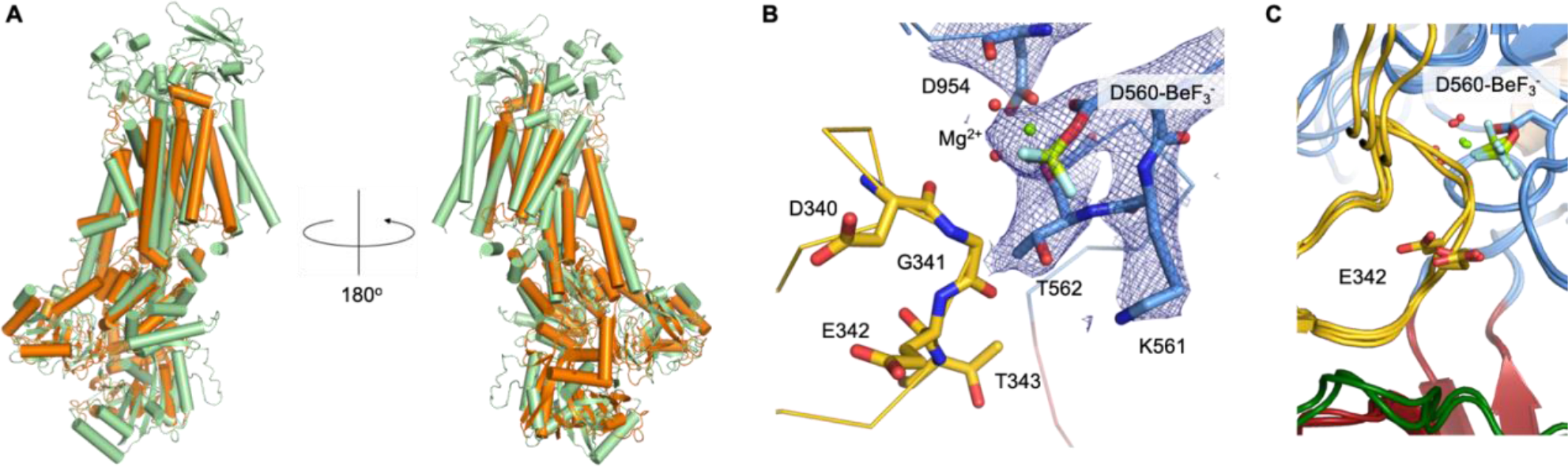
Overall conformation. A) Alignment of Drs2p-Cdc50p E2P^active^ and SERCA (PDB 3B9B^31^). Drs2p-Cdc50p is green, SERCA is orange. (Superposition of C-alpha carbons excluding the N-domain) B) The phosphorylation site of E2P^active^ with density for the BeF_3_^−^ and Mg^2+^ ion and coordinating residues. A characteristic E2P conformation of the dephosphorylation loop with the glutamate pointing away from the phosphorylation site is shown in stick representation. C) The three Drs2p-Cdc50p structures aligned based on the P-domain, with Asp560-BeF_3_^−^ and E342 shown to illustrate the similar conformations. Colors as in Figure 1A.

### Autoinhibition and PI4P binding

We were able to map three distinct intermediate states leading from autoinhibition to activation by variations of sample lipids and inhibitors and using constructs containing intact and truncated C-terminal tails. In E2P^inhib^, the autoinhibitory C-terminus forms an extensive interface that spans 56 residues (residues 1252-1307) along the P- and N-domains, reaching the A-domain (Figure 3A). A short helical segment of the C-terminal tail (H1^C-tail^, residues 1252–1263) interacts with a unique helical insertion on the P-domain, while the rest of the tail extends to interact with the N- and A-domains overlapping with the nucleotide binding site (Figure 3B). Here a conserved GFAFS motif (residues 1274-1278) is positioned at the vertex between the P, N and A domains before extending into the interface of the N- and A-domains (Figure 3A). The autoinhibited state is further stabilised by the clamping of a short loop region in the N-domain (698-704) over the GFAFS motif. Proteolysis of the C-terminus at residue 1290 results in 10-20 fold activation compared to wild-type Drs2p-Cdc50p^25^ and it removes the bulk of the C-terminal residues that interact with the A-domain as well as the 46 unmodeled terminal residues. However, truncation at residue 1302 maintains autoinhibition^25^ and preserves the interactions with the A-domain. This suggests an allosteric mechanism of auto-inhibition, where the cytosolic domains are locked and prevented from undergoing conformational changes. The 16-residue linker between TM10 and the autoregulatory domain is not resolved sufficiently for modelling, but appears at a low density threshold and in 2D class averages (Figure 3C).

**Figure 3:**
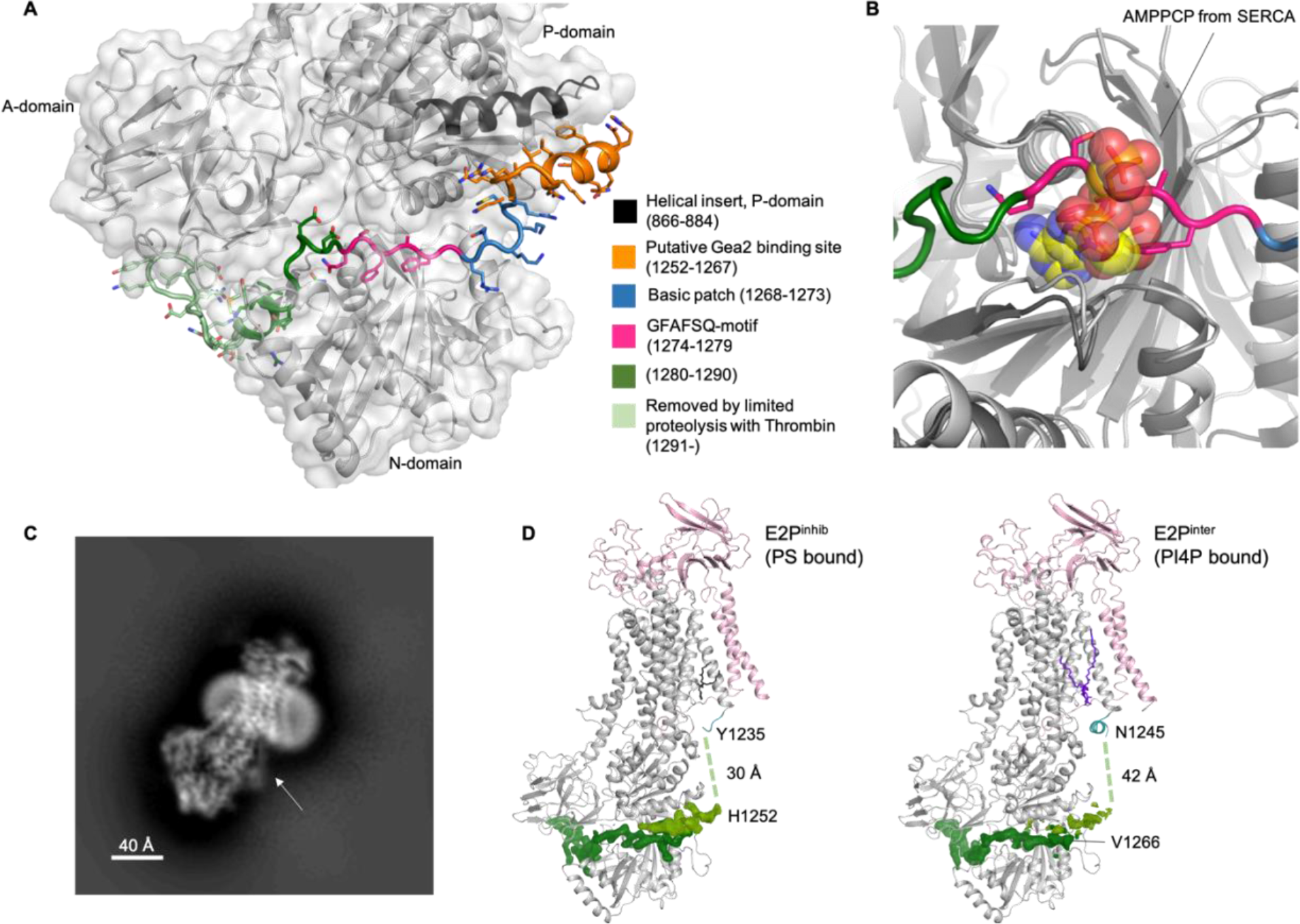
Autoinhibition of Drs2p by its C-terminus. A) The autoinhibitory domain with color-coding of different identified motifs. B) Alignment of N-domains from E2P^inhib^ and AMPPCP-bound SERCA (PDB 1T5S^41^). Drs2p is colored as in A. SERCA is dark grey and AMPPCP is yellow. C) 2D class average from the autoinhibited structure, with an arrow indicating the fuzzy linker between TM10 and the regulatory domain. D) Partial release of autoinhibition upon PI4P binding. The part of the density of the autoinhibitory domain that interacts with the P-domain is shown in lighter green to emphasize its disassociation upon binding of PI4P. The first and last redisues modelled around the disordered linker are listed.

Density is observed for PI4P bound between TM7, 8, and 10 of E2P^inter^ and E2P^active^. Importantly, binding of PI4P is concurrent with formation of an amphipathic helix just after TM10. The amphipathic helix propagates in a direction counter to the remaining part of the auto-inhibitory domain and exerts a mechanical pulling force that dissipates the autoinhibitory interactions of the H1^C-tail^ helical segment with the P-domain. As a result, only downstream interactions of the C-terminus with the N- and A-domain remain intact (Figure 3D). Interestingly, H1^C-tail^ coincides with the previously reported Gea2p binding site, and its release thus mediated by PI4P exposes it for interaction.

The position of the PI4P glycerol backbone is stabilized by interactions with several positively charged residues located in the TM region of Drs2p, but our structures suggest that selectivity for PI4P is driven by the interaction of Tyr1235 and His1236, displayed by the amphipathic helix, to the 4-phosphate of PI4P (Figure 4A and B). This PI4P binding site is notably distinct from a previously proposed patch of basic residues (1268-1273), but it explains why C-terminal truncation at residue 1247 preserves PI4P dependence, cf. E2P^active^ (Supplementary Data Figure 2B), while truncation after residue 1232 leads to an PI4P-independent enzyme^23^.

**Figure 4:**
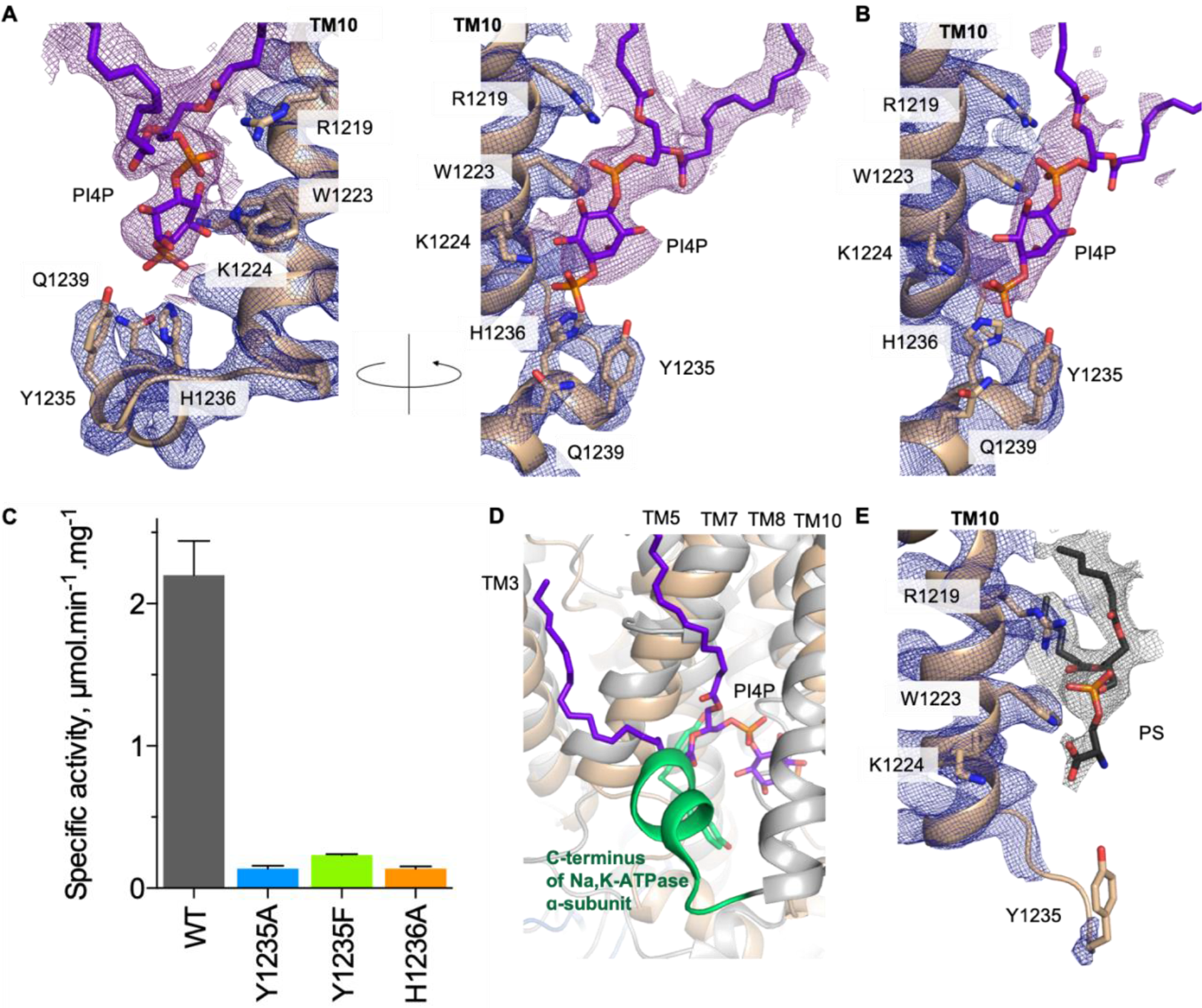
PI4P recognition by Drs2p. A) The PI4P binding site of E2P^active^. The purple density is at a lower threshold (0.75, PyMOL) than the protein (2.5, PyMOL). B) The PI4P binding site of E2P^inter^. The ordered Lys1224 at 3.7Å resolution indicates that it participates in direct contact. The purple and blue densities are at the same threshold (1.5, PyMOL). C) ATPase activity of wild-type and mutants Drs2p-Cdc50p purified in DDM. ATPase activity was plotted as the difference between the rate of ATP hydrolysis observed upon limited proteolysis with trypsin, in the presence of both PI4P and PS, and the rate of ATP hydrolysis observed before adding the various purified protein complexes to the assay medium (in the presence of PS but in the absence of PI4P; see Supplementary Data Figure 5B). Error bars show standard deviation from three different experiments. D) The C-terminus of the Na,K-ATPase (PDB 3KDP^32^), where the terminal 10 residues occupies the same area as PI4P, is shown in green. E) PS binding in E2P^inhib^. Lys1224 is moved away from the binding site, and is not close enough for direct contact. Arg1219 is the only residue still in contact with the glycerophosphate backbone. Grey density is at a lower threshold (1.0, PyMOL), than the blue (2.0, PyMOL).

We further investigated the ability of Tyr1235Ala, Tyr1235Phe, and His1236Ala mutants to hydrolyse ATP. All three mutants were purified with a yield similar to that of the wild-type complex, and Cdc50p interaction remained intact (Supplementary Data Figure 5). The PI4P induced ATPase activity of the Tyr1235Ala, Tyr1235Phe, and His1236Ala mutants however was strongly reduced, less than 10% of wild-type, i.e. consistent with a role in binding PI4P (Figure 5C and Supplementary Data Figure 5B). Owing to the requirement of both PI4P and PS (in addition to the removal of the C-terminal extension of Drs2p by limited proteolysis) for full activation of ATP hydrolysis by Drs2p-Cdc50p, we cannot exclude an effect on PS substrate lipid binding and/or transport. Strikingly, the PI4P binding site of Drs2p is located in the same position as the C-terminal YY motif of the Na,K-ATPase α-subunit^32^, which affects transport function profoundly^33^ (Figure 4D).

**Figure 5:**
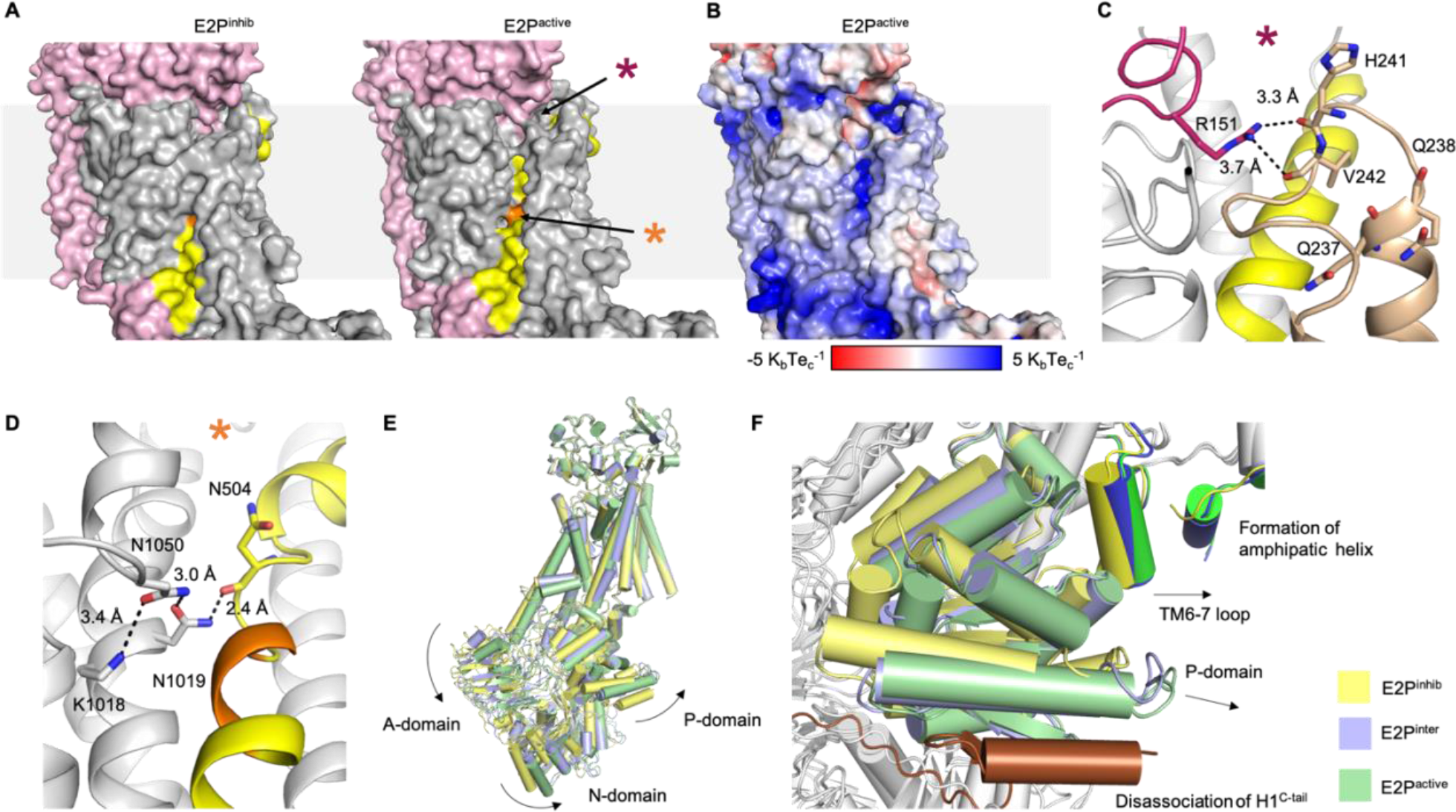
Activation of Drs2p upon PI4P-binding and C-terminal truncation. A) A proposed lipid translocation pathway is exposed upon activation by PI4P binding and C-terminal truncation. TM4 is shown in yellow and the PISL-motif is in orange. E2P^inter^ closely resembles to E2P^inhib^ (not shown). The asterisk marks the location of C) and D). B) Electrostatic surface of the open pathway. Electrostatic surfaces from APBS^42,43^ of all structures are shown in Supplementary Data Figure 5A. C) Interaction between Cdc50p Arg151 and TM1-2 loop of Drs2p in E2P^inhib^. Colors as in A) with TM1-2 in tan. The location within the structure can be seen in A). D) The positive charge of Lys1018 near the PISL-motif of TM4. The location within the structure can be seen in A). E) Alignment of the three Drs2p-Cdc50p structures based on Cdc50p and TM7-10 of Drs2p. E2P^inhib^ is yellow, E2P^inter^ is blue, and E2P^active^ is green. F) View of the P-domains from D with the TM6-7 loop and C-terminus in slightly darker colors and the autoinhibitory domain of E2P^inhib^ in brown.

In E2P^inhib^, the amphipathic helix is not present, but the vacant PI4P site contains a lipid with a significantly smaller head group density with no specific, supporting interactions (Figure 4E). We modelled it as phosphatidylserine, which is the sole lipid added to the sample after extraction from the membrane, but we presume the site accepts regular phospholipids in a nonselective manner in this autoinhibited state.

### A putative substrate entry site

The structures of E2P^active^ and E2P^inter^ are largely similar. However, the lack of an autoinhibitory domain in E2P^active^ allows for slight rearrangements of the N- and A-domains. The movement of the A-domain results in a more open conformation of the TM domain, where TM1 and in particular TM2, extending directly into the A-domain, moves away from the bulk of the TM domain and exposes TM4 to the lumenal leaflet of the membrane. Consistently, the conserved, unwound segment of TM4 (PISL), which is exposed by this movement, has been implicated in lipid transport^14^ (Figure 5A-B). The lumenal opening towards TM4 is lined by TM1, 2, and 6, and we propose that the cleft marks the entry of a substrate lipid transport pathway. The cleft partially overlaps with a previously proposed entry gate and residues important for lipid specificity^12^. In particular Gln237, part of a conserved QQ-motif at the end of TM1, points into the open cleft, supporting its role in substrate specificity (Figure 5C). The cleft is also consistent with the proposed hydrophobic gate model^7^ with a central role for the conserved isoleucine of the PISL motif^14^, although the conformation of TM1-2 in E2P^active^ (and the other E2P sub-states reported here) and earlier homology models are different. Interestingly, while the putative lipid entry pathway is reminiscent of that described for scramblases^10,34^, it does not extend to span both leaflets of the membrane, thus highlighting a fundamental requirement for an alternating access mechanism of lipid movement against its gradient.

Chimera constructs and mutants of the TM1-2 loop in Dnf1p and Drs2p have indicated its role in lipid binding and specificity^12^. A conserved arginine in the ecto-domain of Cdc50p (Arg151) reaches towards the proposed entry site where it could help orient the lumenal loop between TM1 and 2 and guide lipid binding (Figure 5C). Mutations in the TM1-2 loop of Na,K-ATPase confer ouabain resistance^35^ and the cleft coincides with the ouabain binding site in the Na,K-ATPase (Supplementary Data Figure 6B and D). Extension of this cleft towards the cytosolic side overlaps with a lipid binding site and cyclopiazonic acid inhibitory site in SERCA (Supplementary Data Figure 6C and D). Such a proposed exit site to the cytoplasmic leaflet would be expected to emerge only in a subsequent E2-E1 transition of the functional cycle, but the presence of binding pockets and lipid-like ligands in ion pumping P-type ATPase nevertheless hints at possible evolutionary links to the lipid flippases.

**Figure 6:**
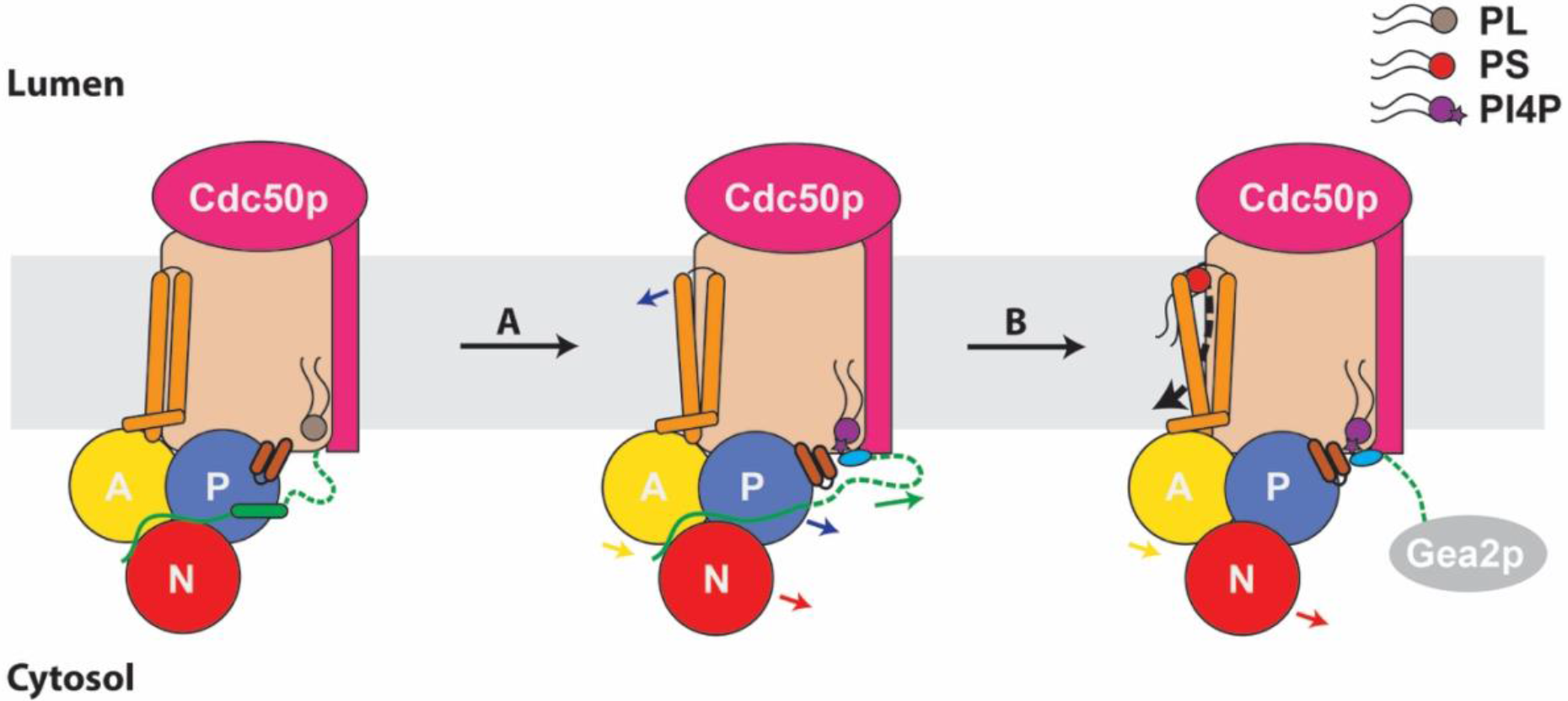
Proposed autoregulation mechanism. Schematic representation of Drs2p-Cdc50p, with functionally important domains and regions colored. Drs2p: A domain (yellow), P-domain (blue), N-domain red, TM1-2 (orange rods), TM6-7 loop (brown rods), TM10 amphipathic helix (light blue), C-terminus (green). Cdc50p (pink). Lipids are identified as per internal legend. The grey ellipse denotes an auxiliary protein e.g., Gea2p that can sequester the C-terminus to prevent rebinding. A) Displacement of the allosteric phospholipid (PL) by PI4P triggers the formation of the amphipathic helix TM10, which disrupts the binding of H1^C-tail^ to the P-domain, resulting in a rigid body movement of the cytosolic domains together with the TM6-7 loop and a slight movement of TM1-2. B) Full displacement of the C-terminus from the cytosolic domains results in a slight movement of the N- and A-domains, which primes the opening of the TM domain by a movement of TM1-2 for substrate lipid binding.

Unlike for the ion transporting P2-ATPases (Supplementary Data Figure 6E and F), negative charges are absent in the core TM structure of Drs2p, but the potentially positively charged Lys1018 of TM5 interacts with Asn1050 at a bulge of TM6. Lys1018 could play a stabilizing, yet dynamic role in the core of the transmembrane domain (Figure 5D). This residue has previously been implicated in transport in bovine ATP8a2^ref 7^, and it projects a potentially positive charge at the middle of the transmembrane pathway (Figure 5A-B,D). The estimated p*K*_a_ of Lys1018 is around 6.8 for E2P^inhib^ and 7.4 for E2P^active, ref. 35,36^indicating that it may switch between a neutral and a positively charged state as part of dynamic interactions with a negatively charged lipid head group.

### Transport mechanism

The respective orientation and positions for the P and N domains of Drs2p are largely invariable in transition from a constrained E2P^inhib^ to the E2P^inter^ and E2P^active^ states. A rigid body movement of those domains relative to the membrane together with a progressive rotation of the A-domain is apparent from an alignment on TM7-10 (Figure 5E). This rigid body movement is also echoed in a concomitant movement of the adjacent TM6-7 loop towards the amphipathic helix formed upon PI4P binding (Figure 5F). Interestingly, the position of the dephosphorylation loop remains constant with respect to the P-domain (Figure 2C). This suggests that binding of PI4P leads to the movement of the P-domain. The subsequent removal of the C-terminus would allow for increased mobility of the A-domain, which would explain why PI4P alone is not sufficient for full activation of the intact enzyme.

Taken together, we propose the following model for auto-regulation and exposure of the lipid entry site. E2P^inhib^ has a closed TM domain with the cytosolic domains locked by the autoinhibitory C-terminus (Figure 6). PI4P binding to the TM domain promotes the folding of an amphipathic helix right on the C-terminal side of TM10. Formation of this helix has two effects: 1) it causes the remainder of the C-terminus to partially unfold, thus destabilizing its interaction with the P-domain through the displacement of H1^C-tail^ with the putative Gea2p binding site; and 2) it forms an interaction site for the TM6-7 loop to form, which then moves with the P-domain. Together these two movements shift the P and N domains towards the amphipathic helix largely as a rigid body. A progressive rotation of the A domain leads to a subtle movement of TM2 and to a lesser extent TM1 giving the E2P^inter^ state. Full displacement of the C-terminus leads to the E2P^active^ state where a further rotation of the N- and A-domains drives the opening of a putative substrate binding site through the movement of TM2 away from TM6. We anticipate that subsequent PS (or PE) substrate lipid binding will be associated with further conformational changes of the TM domain, as the enzyme dephosphorylates.

## Conclusion

The structures of Drs2p-Cdc50p presented here provide the first insights into the architecture of the P4-ATPase lipid flippases and the location of determinants of lipid and Cdc50p specificity. The structure of the autoinhibitory domain and the mechanism for relief of autoinhibition through a regulatory PI4P site explain details of regulation of P4-ATPases that may also be targeted for specific modulation of lipid flippase activity. Further structures capturing progressive states of substrate lipid binding, dephosphorylation, lipid translocation and cytoplasmic release will be required to further elucidate the lipid flippase mechanism.

## Acknowledgements

The authors wish to thank J. Karlsen for scientific computing and molecular graphics facilities and assistance to their use, A. M. Nielsen and T. Klymchuk for technical assistance, and C. Grønberg for early contributions on sample preparation. We thank D. Mills and the staff at the Department of Structural Biology (MPI of Biophysics, Frankfurt/Main) for support on data collection, discussions and support. Support for this work was provided by grants from the Danish National Research Foundation for the PUMPkin center of excellence, and from the Lundbeck Foundation for the Brainstruc center of excellence (2015–2666) to P.N., by an EMBO Long-Term Fellowship to M.R.A., by postdoctoral grants from the Danish Research Council and the Lundbeck Foundation to J.A.L., by a PhD fellowship from the Boehringer-Ingelheim Fonds to M.T. (more….)

## Author Contributions

P.N. and G.L. conceived the project, and J.A.L., T.H.B. and P.N. defined the cryo-EM study with A.M. and W.K. The samples were characterized and developed by M.T., J.J.U., J.A.L., and M.R.A., and exploratory electron microscopy studies were performed by J.A.L., M.T., J.J.U. and T.H.B. Cryo-EM analysis was performed by M.T., D.J., J.A.L., T.H.B., and A.M. Data processing and 3D reconstruction was performed by M.T. with support and advice from D.J., J.A.L., and A.M. Model building and refinement was performed by M.T. and J.A.L, with assistance from J.J.U. Mutant forms were prepared and functionally characterized by T.D., C.M. and G.L. P.N. and J.A.L. supervised the project together with A.M. The manuscript was drafted by M.T., J.A.L. and P.N. All authors commented on the manuscript.

## Author Information

Cryo-EM maps for the *S. cerevisiae* Drs2p-Cdc50p (UniProt ID XXXXX) in the E2–BeF3 (E2inhib), E2-BeF3-PI4P (E2Pinter), and C-terminally truncated E2-BeF3-PI4P (E2Pactive) forms are available on the xxxxx server, and coordinates of the atomic structures have been deposited in the Protein Data Bank (PDB) under accession numbers XXXX, YYYY, and ZZZZ. Reprints and permissions information is available at www.nature.com/reprints. The authors declare no competing financial interests. Readers are welcome to comment on the online version of the paper. Correspondence and requests for materials should be addressed to P.N, A.M., or G.L.

---

**Supplementary Data Figure 1:**
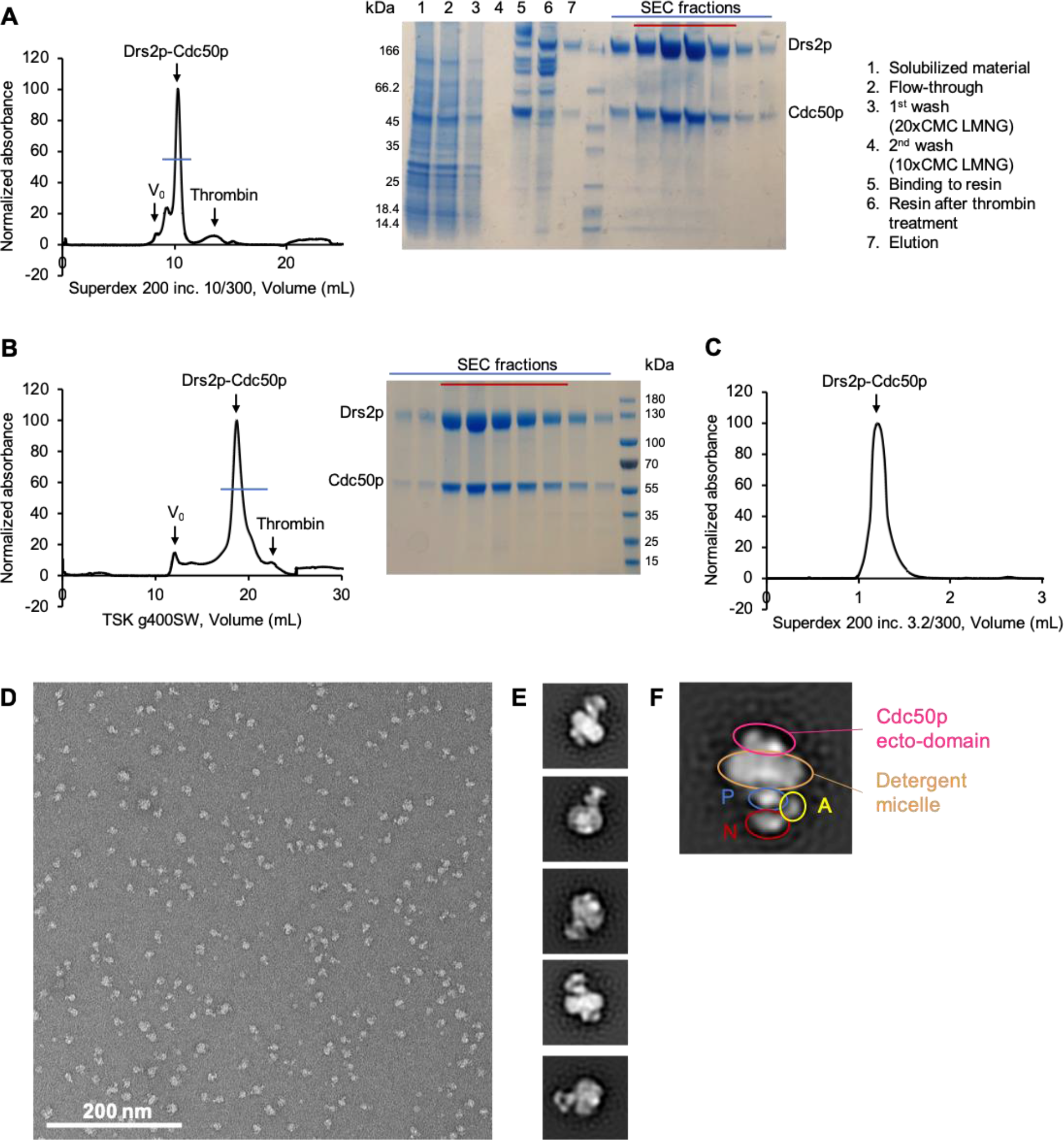
Purification and negative stain electron microscopy of samples for structural studies. A) Chromatogram and gel of Drs2p ΔN104/Cdc50p purification. Red line shows the pooled fractions. B) 1st SEC of Drs2p ΔNC/Cdc50p with gels showing fractions. Red line shows the pooled fractions. C) 2nd SEC of ΔNC, used for freezing grids. D) A representative negative stained micrograph of autoinhibited Drs2p ΔN104/Cdc50p in LMNG. E) Representative 2D class averages of the sample in D) shows well defined and homogeneous particles with recognizable P-type ATPase features, showing that the sample is suitable for further study by cryo-EM. F) Enlarged 2D class average with highlight of the recognizable domains of Drs2p-Cdc50p.

**Supplementary Data Figure 2:**
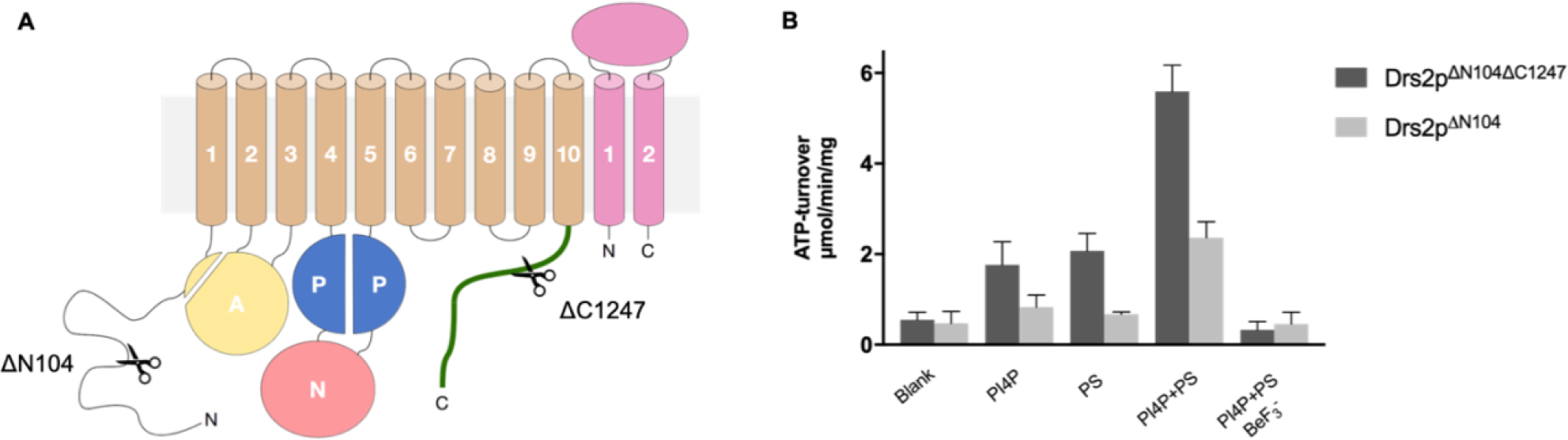
Activity of samples used for structural studies. A) Topology of Drs2p-Cdc50p with indication of the cleavages at the termini of Drs2p of the constructs used for structural studies (ΔN104: all constructs, ΔC1247: E2P^active^). Cdc50p is pink, while for Drs2p the TM-domain is tan, the A-domain is yellow, the P-domain is blue, the N-domain is red and the autoinhibitory C-terminus is green. B) Specific activity of Drs2p ΔNC/Cdc50p and Drs2p ΔN104/Cdc50p in LMNG measured by the Baginski Assay. Where present, PS C(8:0), Brain PI4P and BeF_3_^−^ were added to final concentrations of 78μg/mL, 20μg/mL and 5mM, respectively.

**Supplementary Data Figure 3:**
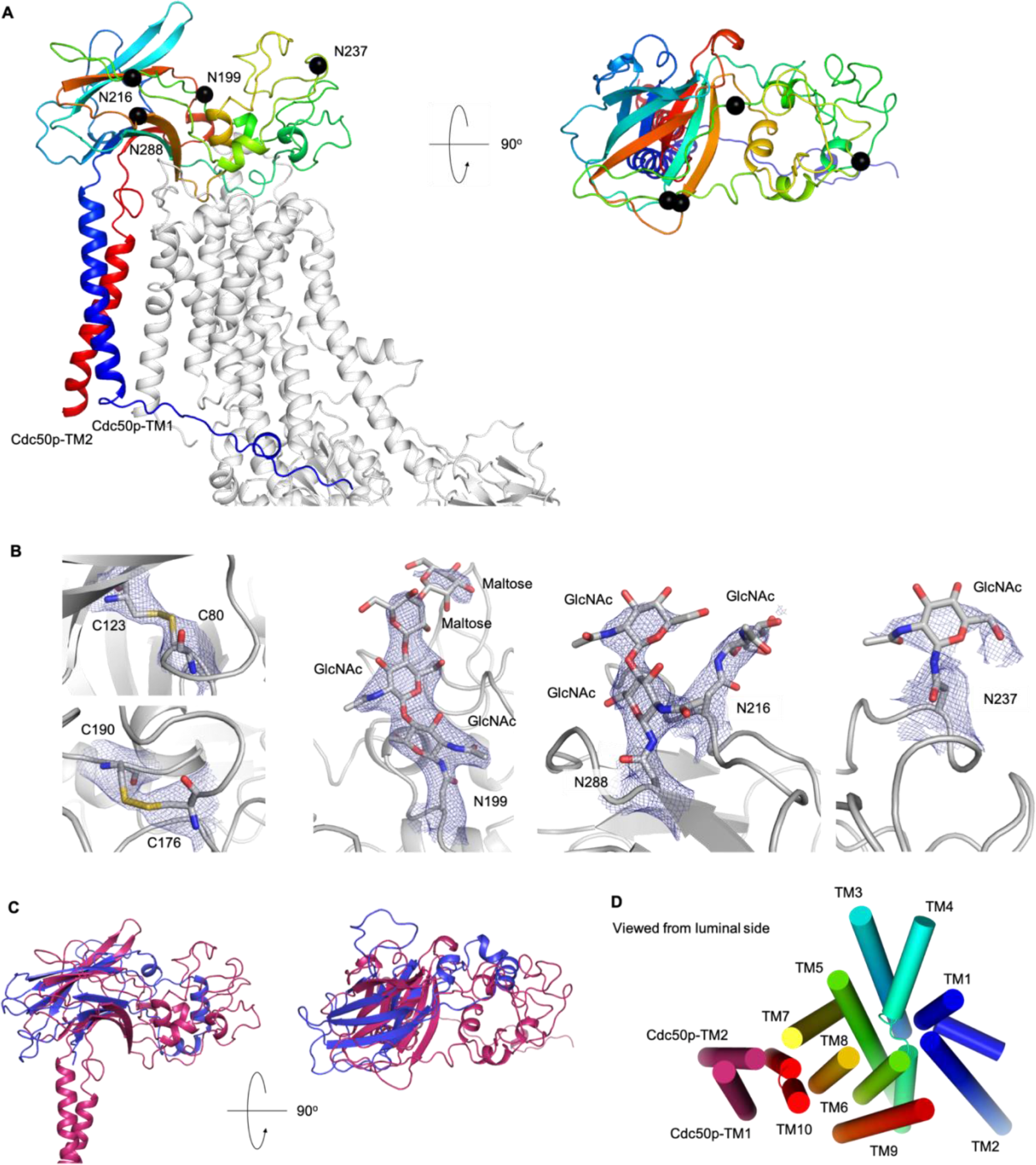
Fold of Cdc50p. A) Cdc50p from E2P^inhib^ colored by rainbow. C_α_-carbons of asparagines carrying glycosylations are shown as black spheres. Drs2p is shown in grey. B) Disulfides and glycosylations of Cdc50p from E2P^inhib^. Cys176-Cys190 and Asn237-GlcNAc are shown at lower thresholds than the rest. C) Alignment of Cdc50p and a monomer of human seipin (PDB 6DS5^44^), a lipid binding protein illustrates the similar folds, although loops of Cdc50p are more extensive. The sequence identity between the two proteins is only 4%. Transmembrane helices of seipin extending from similar positions as the helices of Cdc50p are missing in the structure, but may extend into the membrane in a similar way as observed for Cdc50p. D) The TM-helices of the complex viewed from the luminal side. The helices of Drs2p are colored in rainbow.

**Supplementary Data Figure 4:**
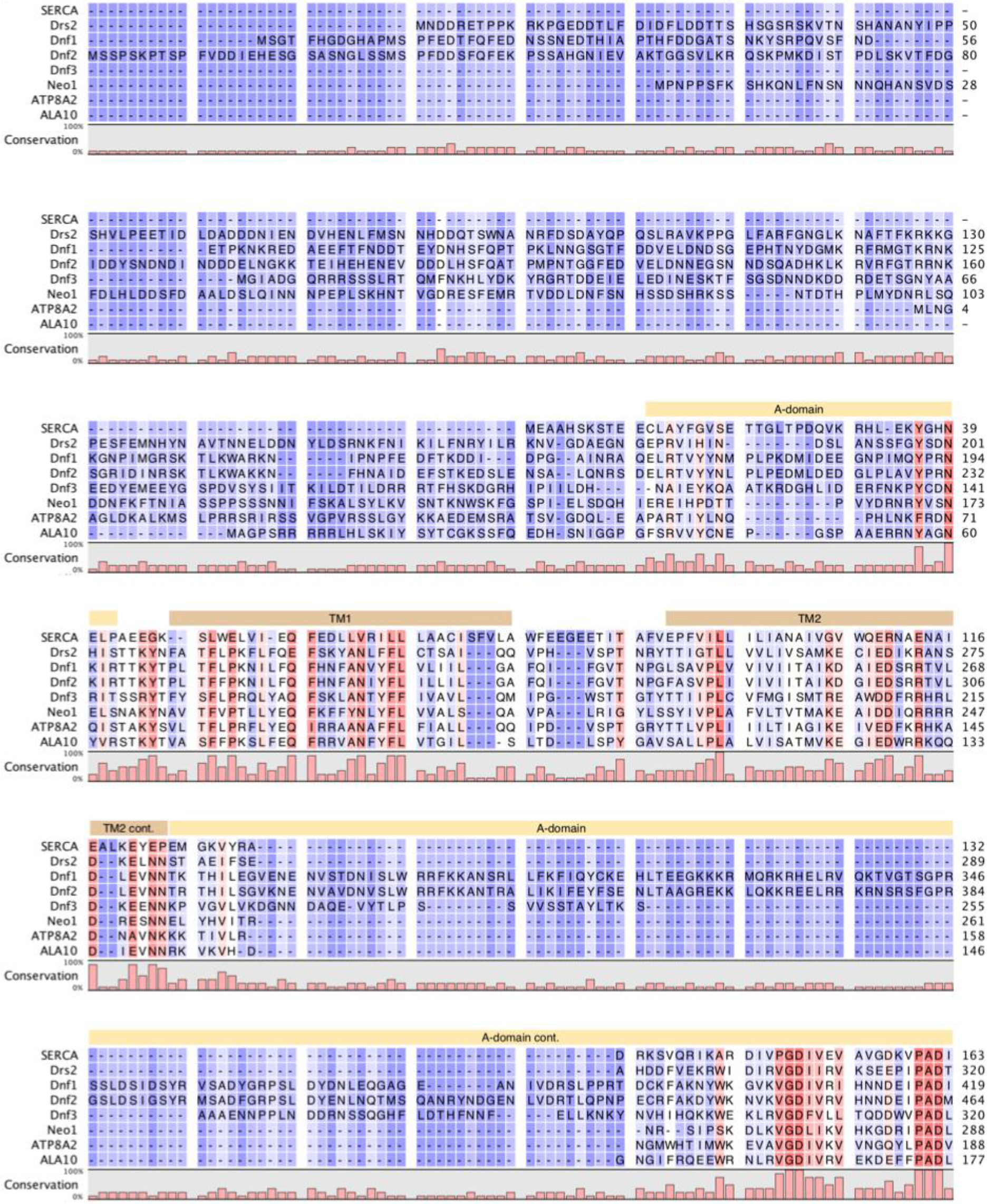
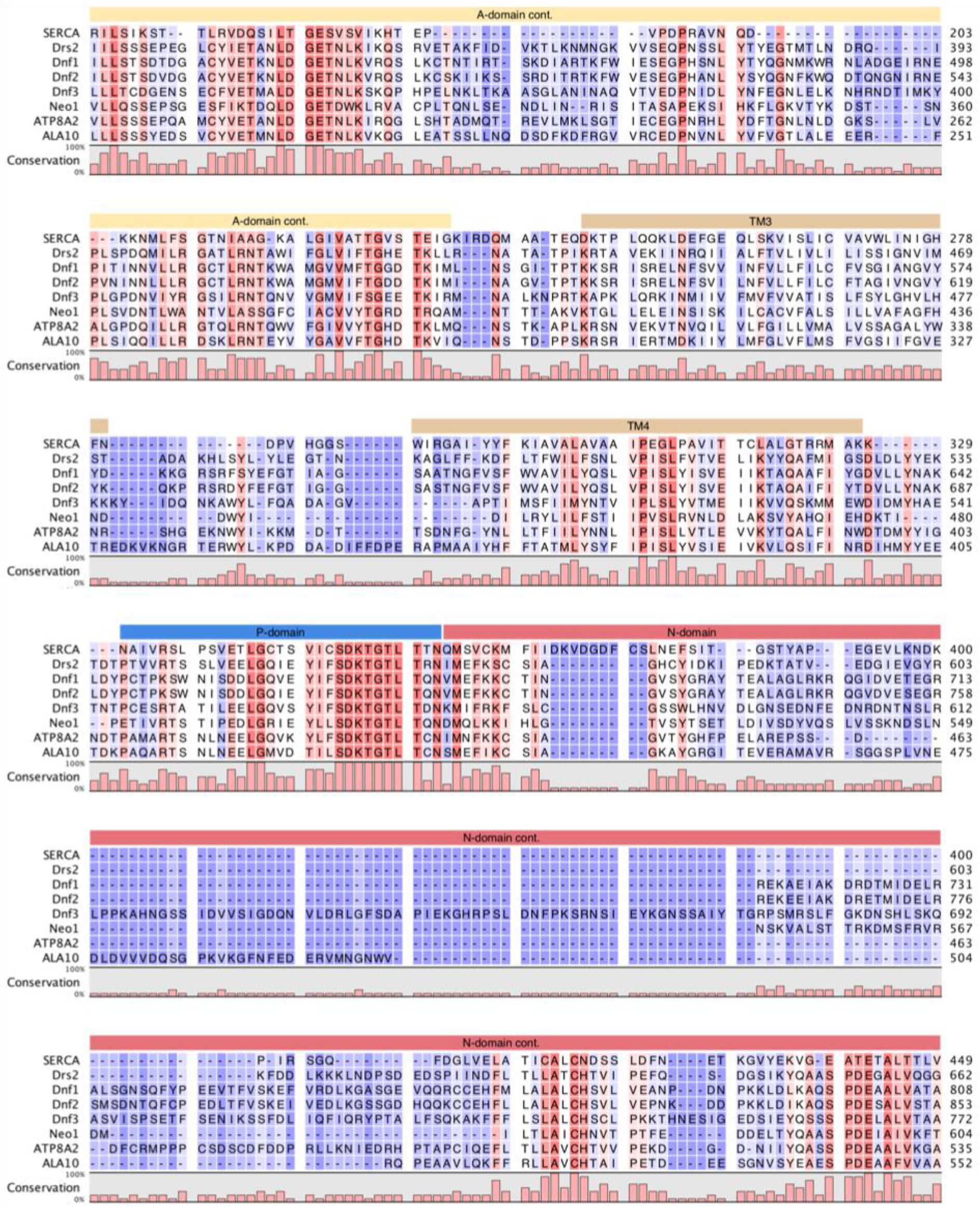
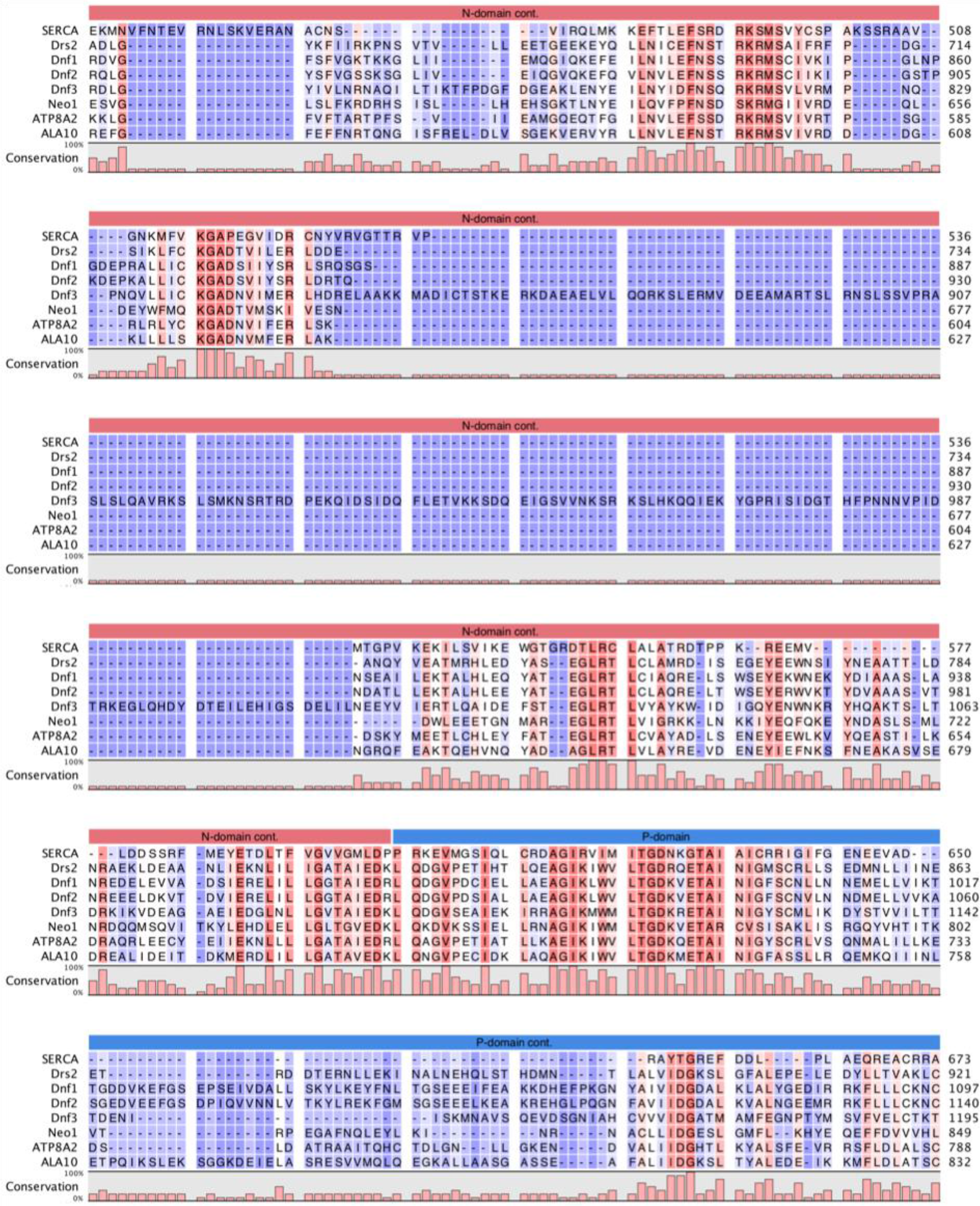
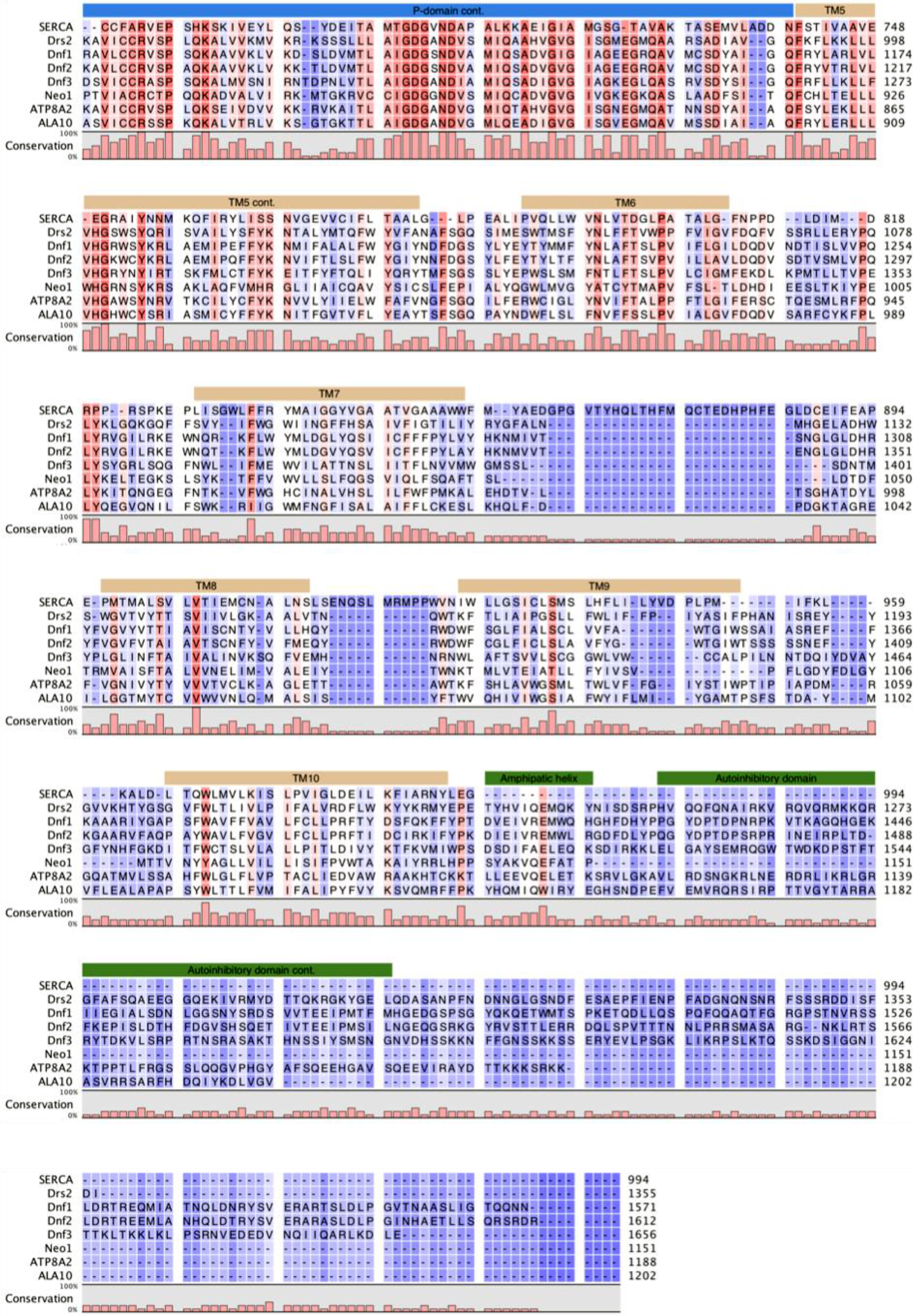
Sequence alignment of select P4 ATPases and structural alignment with SERCA. Sequence alignment of the five *S. cerevisiae* P4 ATPases, the human ATP8A2, and *A. thaliana* ALA10, aligned by Clustal Omega^39,40^, along with a structural alignment between Drs2p from E2Pactive and SERCA-BeF_3_^−^ (PDB 3B9B) performed by the Dali Server’s pairwise alignment^45^ separately for the N-domain of Drs2p and for the rest of the protein, as the N-domain is very flexible and can distort the alignment. The structural alignment was adjusted to the full-length Drs2p-sequence. For the rabbit SERCA it only includes the sequence of the structure that is shown, meaning that the C-terminal 7 residues are missing. The shading indicates conservation (blue 0% – red 100%). Above the sequences the domains and transmembrane helices of Drs2p are indicated in the same color scheme as Figure 1A. Uniprot identifiers: SERCA – P04191, Drs2p – P39524, Dnf1p – P32660, Dnf2p – Q12675, Dnf3p – Q12674, Neo1p – P40527, ATP8A2 – Q9NTI2, ALA10 – Q9LI83.

**Supplementary Data Figure 5:**
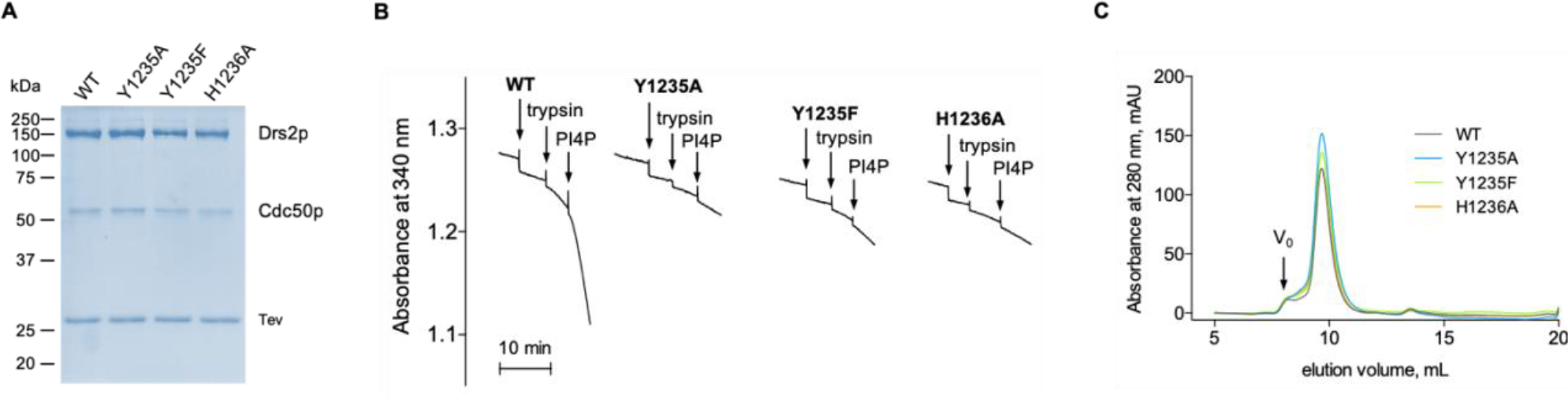
Characterization of PI4P-binding mutants. A) Coomassie-blue stained SDS-PAGE of streptavidin-purified wild-type and mutant Drs2p-Cdc50p. Tobacco Etch Virus (TEV) protease is used to release the complex from streptavidin beads. B) ATPase activity of PI4P-binding mutants, using an enzyme-coupled assay. The assay medium contained 1 mM ATP, 0.1 mg.mL^−1^ POPS and 1 mg.mL^−1^ DDM in SSR buffer. The rate of ATP hydrolysis was continuously recorded at 340 nm upon subsequent addition of 2 μg.mL^−1^ of the purified complex, 5 μg.mL^−1^ trypsin, and 0.025 mg.mL^−1^ PI4P.C) Size-exclusion chromatography on a Superdex 200 10/300GL column. The arrow indicates the dead volume (V_0_).

**Supplementary Data Figure 6:**
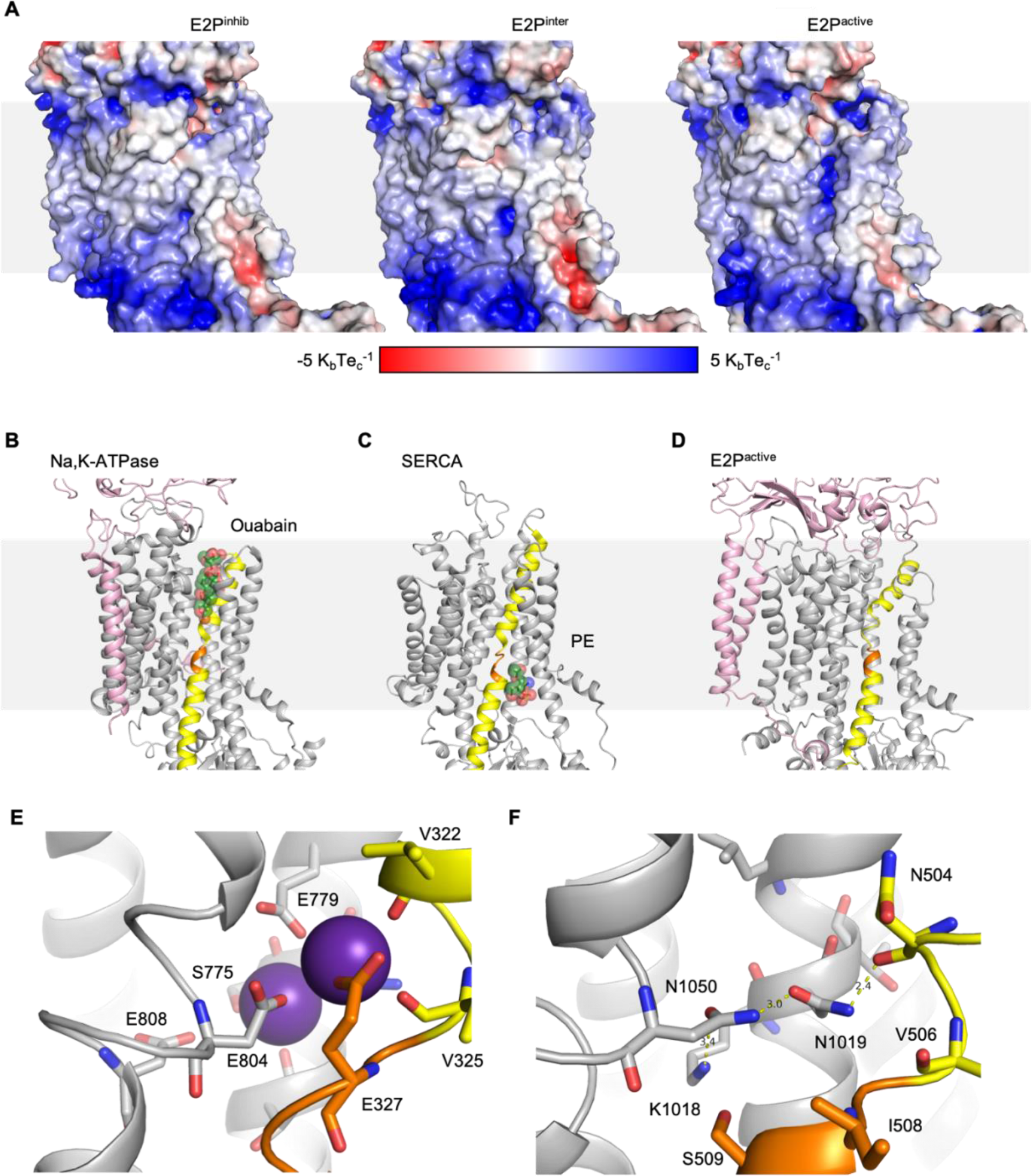
Electrostatics of proposed lipid translocation pathway. A) A proposed lipid translocation pathway is exposed upon activation by PI4P binding and C-terminal truncation. Electrostatic surfaces from APBS^42,43^. Complement to Figure 5A. B) The E2P-ouabain complex structure of Na,K-ATPase (PDB 4HYT). The β- and ɣ-subunit are pink, TM4 is yellow, the PEGL-motif is orange and the ouabain ligand primarily green. Note the clear overlap with the opening in Drs2p-Cdc50p occurring upon activation, Figure 5A. C) An E2 state of SERCA with a PE molecule bound between TM2 and 4 (PDB 2AGV^46^). Colors as (B). D) E2P^active^ in the same orientation and with colors as B-C. E) Potassium binding in Na,K-ATPase (PDB 3KDP^32^), showing the coordination of ions in site I and site II by negatively charged and polar residues. TM4 is colored yellow with the PEGL-motif in orange, and the bound potassium as purple spheres. F) Sites and residues corresponding to the ion binding sites of Na,K-ATPase (PDB 3KDP) shown in (D) from E2P^active^. TM4 is yellow with the PISL-motif in orange, and stabilizing hydrogen bonds are shown.

**Supplementary Data Figure 7:**
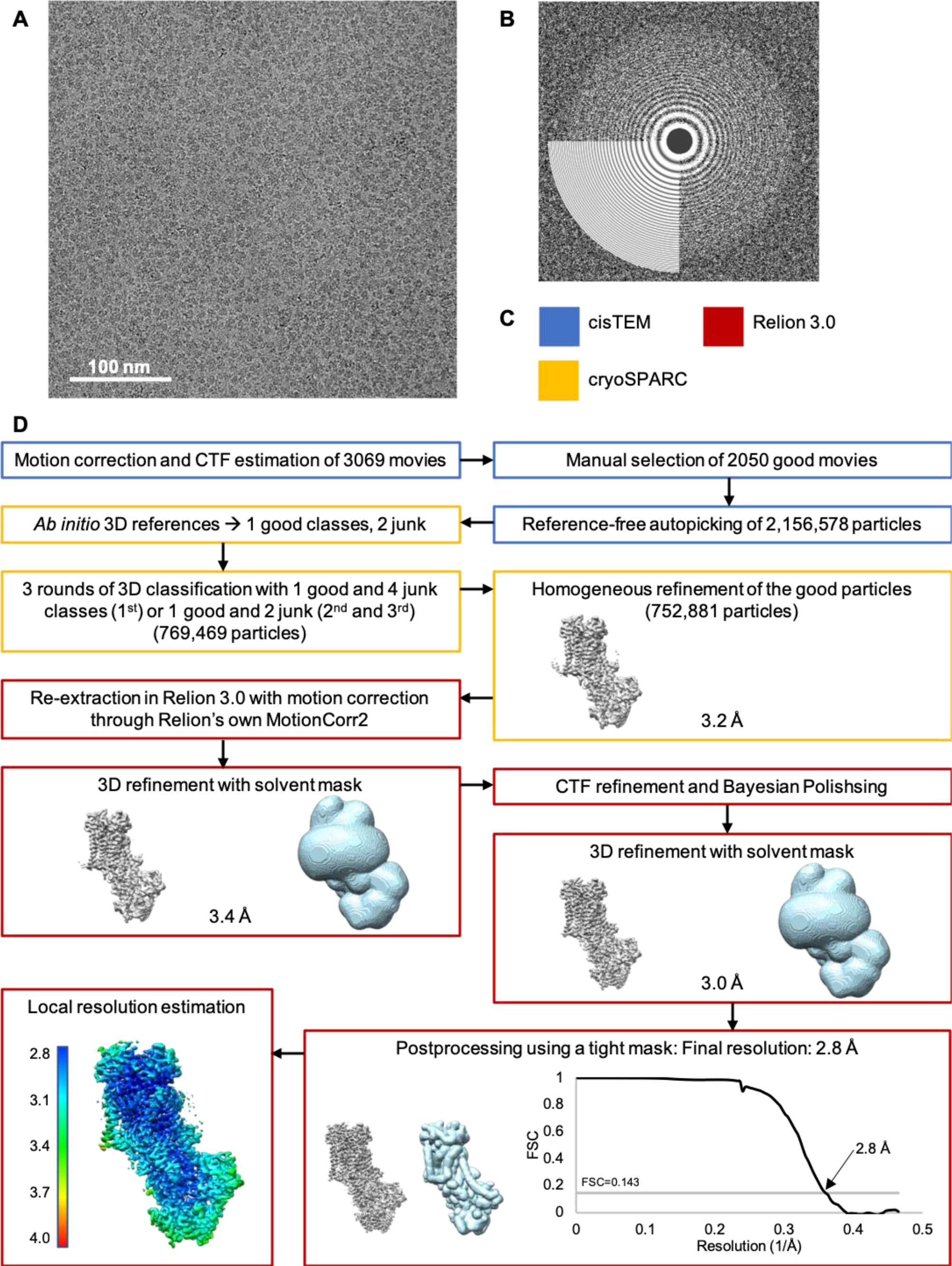
Processing pipeline for cryo-EM data of Drs2p-Cdc50p E2P^inhib^. A) Representative motion-corrected and dose-weighted micrograph (defocus 1.6 μm) of autoinhibited Drs2p ΔN104-Cdc50p in LMNG, frozen at a concentration of 0.6 mg/ml. B) Fourier power spectrum of the micrograph shown in (A), with fit from CTFFIND 4.1 through cisTEM, which extends to 3 Å. C) Color code of processing software. D) Data processing workflow with indication of the number of particles remaining after each step at which particles were discarded. The densities resulting from 3D refinement are shown in grey, while relevant masks are shown in light blue. The resolutions listed for 3D refinements are at FSC=0.143.

**Supplementary Data Figure 8:**
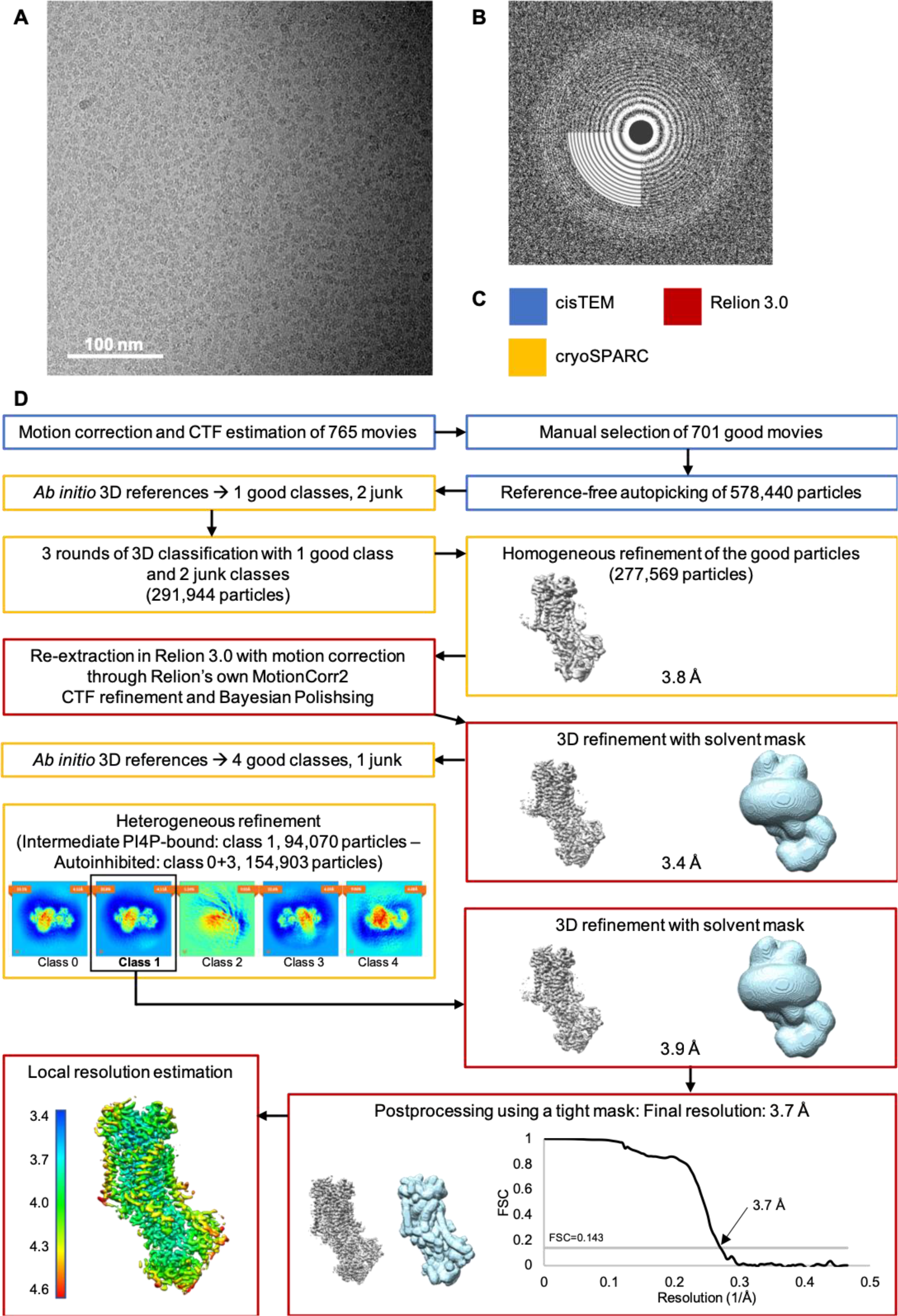
Processing pipeline for cryo-EM data of Drs2p-Cdc50p E2P^inter^. A) Representative motion-corrected and dose-weighted micrograph (defocus 1.5 μm) of Drs2p ΔN104-Cdc50p with an intact C-terminus in LMNG, frozen at a concentration of 0.6 mg/ml in the presence of 75 μg/ml Brain PI4P. B) Fourier power spectrum of micrograph in (A), with fit from CTFFIND 4.1 through cisTEM, which extends to 5 Å. C) Color code of processing software. D) Data processing workflow with indication of the number of particles remaining after each step at which particles were discarded. Densities resulting from 3D refinement are shown in grey, while relevant masks are shown in light blue. The resolutions listed for 3D refinements are at FSC=0.143. The 3.4 Å-refinement indicated a mixed state around TM10. To classify structural heterogeneity due to incomplete binding of PI4P, new ab initio references were generated in cryoSPARC allowing for high similarity, as the conformations were expected to be similar. Two different conformations resulted: the autoinhibited one and a PI4P-bound. The autoinhibited was identical to E2P^inhib^ and adding these particles to that dataset did not improve the reconstruction. The PI4P-bound conformation was further refined in Relion.

**Supplementary Data Figure 9:**
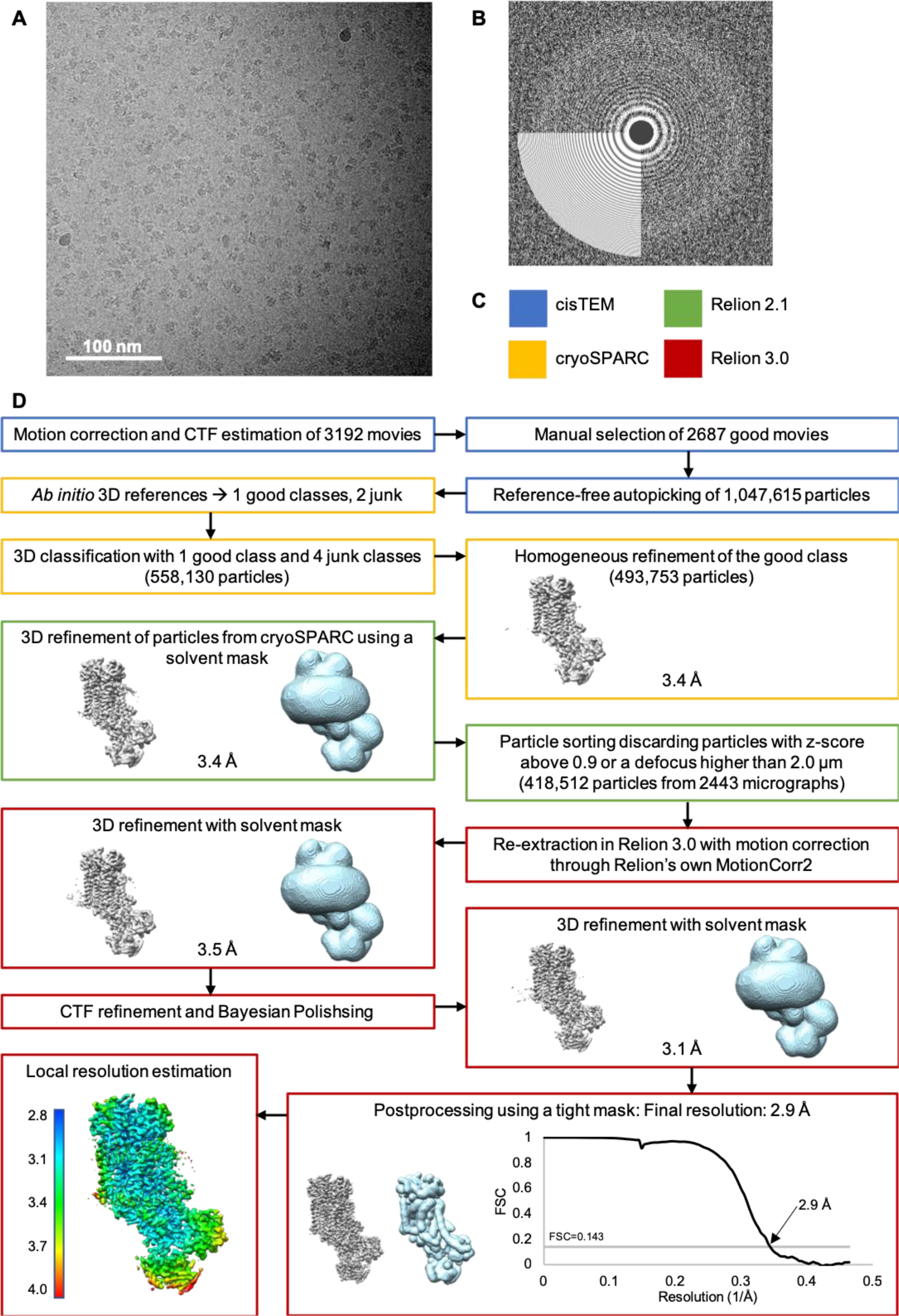
Processing pipeline for cryo-EM data of Drs2p-Cdc50p E2P^active^. A) Representative motion corrected and dose weighted micrograph (defocus of 1.7 μm) of C-terminally truncated Drs2p ΔNC-Cdc50p in LMNG, frozen at a concentration of 0.6 mg/mL in the presence of 75 μg/mL Brain PI4P. B) Fourier power spectrum of the micrograph shown in (A), as well as the fit from CTFFIND 4.1 through cisTEM, which extends to 3 Å. C) Color code of processing software. D) Data processing workflow with indication of the number of particles remaining after each step at which particles were discarded. Densities resulting from 3D refinement are shown in grey, while relevant masks are shown in light blue. The resolutions listed for 3D refinements are at FSC=0.143.

**Supplementary Data Figure 10:**
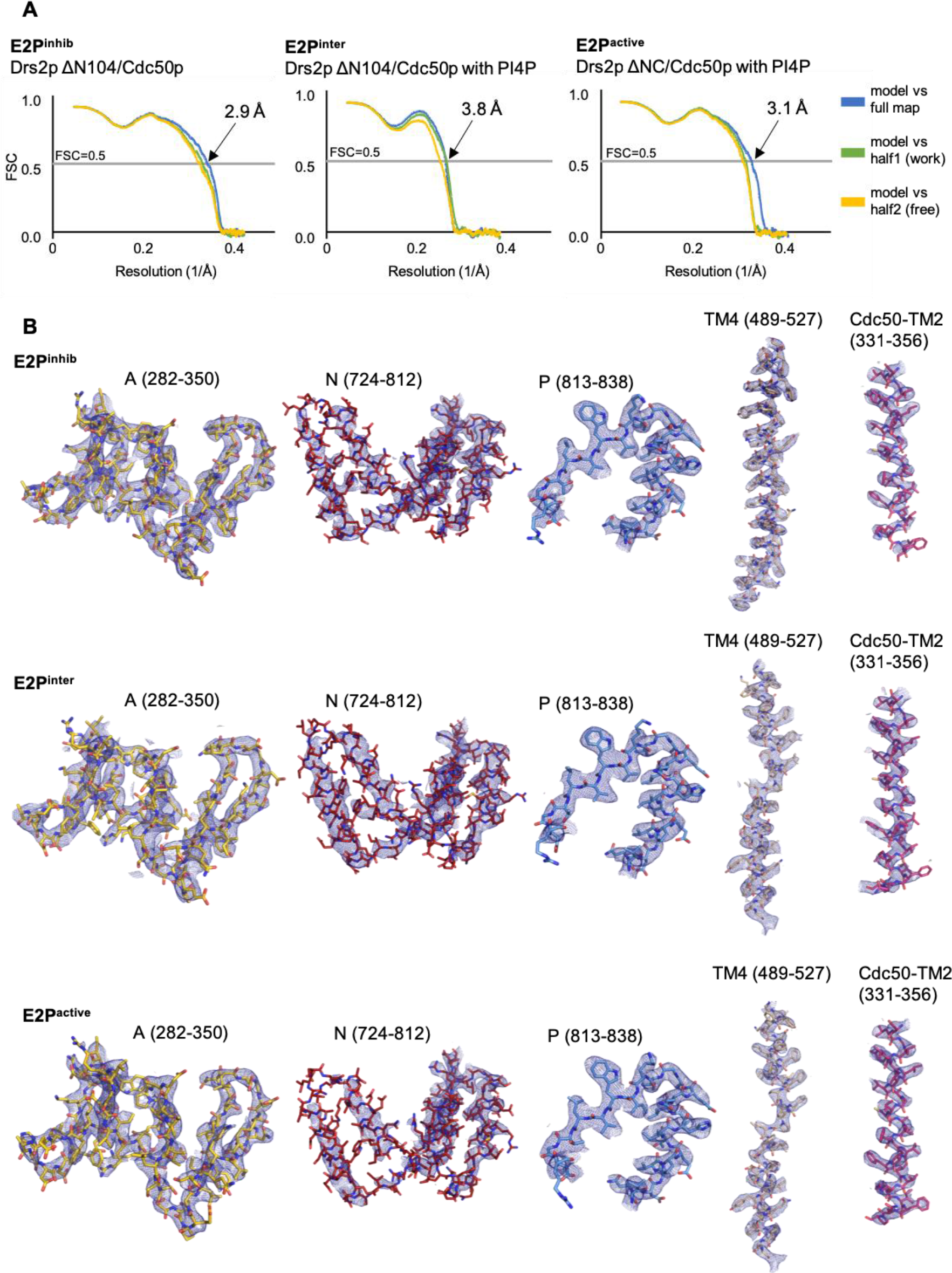
Model validation and representative densities. A) Cross-validation FSC curves for map-to-model fit produced by Mtriage^47^. Curves representing model vs. full map are calculated based on the final model and the full, filtered and sharpened map that it was refined against. For the model vs. half-maps, the model (before the final refinement against) was refined against half-map 1 filtered and sharpened as the full map, and FSC-curves were calculated using this refined model against each half-map. B) Representative densities from different areas of the three LocScale-maps. Each segment is labelled with which residues are shown, and demonstrates the quality of the map in specific areas. All are shown at a 1.5 threshold (PyMOL).

**Supplementary Data Table 1:** Cryo-EM data collection, refinement and validation statistics

|  | Drs2p ΔN104/Cdc50p<br>(autoinhibited)<br>(EMDB-xxxx)<br>(PDB xxxx) | Drs2p ΔN104/Cdc50p<br>with PI4P<br>(intermediate)<br>(EMDB-xxxx)<br>(PDB xxxx) | Drs2p ΔNC/Cdc50p<br>with PI4P<br>(activated)<br>(EMDB-xxxx)<br>(PDB xxxx) |
| --- | --- | --- | --- |
| **Data collection and processing** |  |  |  |
| Magnification | 130,000 | 130,000 | 130,000 |
| Voltage (kV) | 300 | 300 | 300 |
| Microscope | Titan Krios | Titan Krios | Titan Krios |
| Electron exposure (e<sup>-</sup>/Å<sup>2</sup>) | 60 | 60 | 56 |
| Defocus range (μm) | 0.3-3.0 (0.4-2.5)* | 0.4-2.4 (0.5-2.4)* | 0.3-3.0 (0.4-2.0)* |
| Pixel size (Å) | 1.077 | 1.077 | 1.077 |
| Camera | Gatan K2 | Gatan K2 | Gatan K2 |
| Number of movies | 3069 (2050)* | 765 (701)* | 3192 (2443)* |
| Symmetry imposed | C1 | C1 | C1 |
| Initial particle images (no.) | 2,157,578 | 578,440 | 1,047,615 |
| Final particle images (no.) | 752,881 | 78,981 | 418,512 |
| Map resolution (Å) | 2.8 | 3.7 | 2.9 |
| FSC threshold | 0.143 | 0.143 | 0.143 |
| Map resolution range (Å) | 2.7-3.4 | 3.6-4.4 | 2.9-4.1 |
| **Refinement** |  |  |  |
| Initial model used (PDB code) | N/A | N/A | N/A |
| Model resolution (Å) | 2.9 | 3.8 | 3.1 |
| FSC threshold | 0.5 | 0.5 | 0.5 |
| Model resolution range (Å) |  |  |  |
| Map sharpening *B* factor (Å<sup>2</sup>)<sup>#</sup> | -40 (-76) | -60 (-115) | -35 (-69) |
| Model composition |  |  |  |
| Non-hydrogen atoms | 11772 | 11791 | 11479 |
| Protein residues | 1449 | 1447 | 1404 |
| Ligands | 2 H<sub>2</sub>O, 1 Mg<sup>2+</sup>, 1 PS,<br>6 NAG, 2 BMA | 2 H<sub>2</sub>O, 1 Mg<sup>2+</sup>, 1 PI4P,<br>6 NAG, 2 BMA | 2 H<sub>2</sub>O, 1 Mg<sup>2+</sup>, 1 PI4P,<br>7 NAG, 4 BMA |
| *B* factors (Å<sup>2</sup>) |  |  |  |
| Protein | 33.75/150.55/73.76 | 35.45/170.28/89.45 | 34.18/187.73/77.30 |
| Ligand | 55.17/99.22/78.05 | 68.87/120.20/92.11 | 51.25/111.21/78.04 |
| Water | 55.49/55.73/55.61 | 77.61/80.06/78.84 | 60.71/61.39/61.05 |
| R.m.s. deviations |  |  |  |
| Bond lengths (Å) | 0.006 | 0.008 | 0.006 |
| Bond angles (°) | 0.868 | 0.995 | 0.932 |
| Validation |  |  |  |
| MolProbity score | 1.40 | 1.61 | 1.36 |
| Clashscore | 3.99 | 5.05 | 3.57 |
| Poor rotamers (%) | 0.23 | 0.23 | 0.16 |
| EMRinger | 3.80 | 2.13 | 3.50 |
| Ramachandran plot |  |  |  |
| Favored (%) | 96.60 | 94.99 | 96.64 |
| Allowed (%) | 3.40 | 5.01 | 3.36 |
| Disallowed (%) | 0.0 | 0.0 | 0.0 |

\* number in parenthesis corresponds to the final range/number.
<sup>#</sup> The B-factor listed is the one used for refinement, while the one in the parentheses is the B-factor estimated by Relion PostProcessing.

## Supplementary Information

### Methods

#### Enzyme-coupled assay

Expression and streptavidin purification of wild-type Drs2p-Cdc50p and mutants for functional studies were carried out as described previously^20,48^. Specifically, we used for this purpose a C-terminal Tobacco Etch Virus (TEV)-cleavable BAD tag and DDM was used throughout the purification procedure. Size-exclusion chromatography was performed on a Superdex 200 Increase 10/300GL column, with a mobile phase containing 0.5 mg.mL^−1^ DDM and 0.025 mg.mL^−1^ POPS. The rate of ATP hydrolysis by PI4P-binding mutants was measured at 30 °C using an enzyme-coupled assay, by continuously monitoring the rate of NADH oxidation at 340nm^49^. The purified Drs2p-Cdc50p complexes were added at about 2 μg.mL^−1^, in a cuvette containing SSR buffer supplemented with 1 mM ATP, 1 mM phosphoenolpyruvate, 0.4 mg.mL^−1^ pyruvate kinase, 0.1 mg.mL^−1^ lactate dehydrogenase, 0.25 mM NADH, 1 mM NaN3, 1 mg.mL^−1^ DDM, and 0.1 mg.mL^−1^ POPS. Trypsin and PI4P were subsequently added to concentrations of 0.05 mg.mL^−1^ and 0.025 mg.mL^−1^, respectively. Conversion of NADH oxidation rates expressed in AU/s to ATPase activities in μmol.min^−1^.mg^−1^ was based on the extinction coefficient of NADH (~6200 M^−1^.cm^−1^) and on Coomassie-Blue stained SDS-PAGE-based quantification of Drs2p using know amounts of SEC-purified Drs2p as standards.

#### Expression and purification of Drs2p-Cdc50p for structural studies

Protein expression in *S. cerevisiae*, membrane harvesting and solubilization were performed as previously described^20,25,48^. Two different preparations were used Drs2p ΔN104/Cdc50p, where Drs2p is missing the first 104 residues, and Drs2p ΔNC/Cdc50p, where residues 105-1247 of Drs2p are present.

#### Affinity chromatography on streptavidin resin and detergent exchange

The BAD-tagged protein was batch bound to free streptavidin sepharose resin (typically 1mL resin per 60mL of solubilized material) for 1 hour at 4^◦^C. Detergent exchange into lauryl maltose neopentyl glycol (LMNG) was performed, by washing with 2 column volumes (CV) SSR with 1mM DTT and 0.2 mg/mL LMNG, followed by washing with 10 CV SSR with 1 mM DTT and 0.1 mg/mL LMNG. The resin was resuspended in 1 CV SSR with 1mM DTT, 0.1 mg/mL LMNG and 50 μg/mL Brain PS (Avanti Polar Lipids). 4 units/mL resin of bovine Thrombin (Calbiochem) was added to cleave the protein off the resin during an overnight incubation at 4^◦^C. The protein was eluted from the resin in 10-20 CV of SSR with 1mM DTT and 0.1mg/mL LMNG, and concentrated to 0.5-1mL in a 100 kDa centrifugal concentrator (Vivaspin) with the sample typically reaching concentrations of 5-10 mg/mL.

#### Cleavage of double truncated construct

To produce the double-truncated Drs2p ΔNC/Cdc50p protein, after elution from the resin and concentration, 4U bovine thrombin per mL of resin used for the purification was added along with 0.025 mg/mL Brain PI4P (Avanti Polar Lipids), followed by incubation at room temperature for 1 hour, before quenching of the protease activity with 1mM PMSF.

#### Size exclusion chromatography

For Drs2p ΔN104/Cdc50p, size exclusion chromatography (SEC) was run on a Superdex 200 Increase 10/300 column on an ÄKTA purifier system at 4°C resulted in a Drs2p/Cdc50p concentration of 0.6 g/mL, which was used directly for in SSR with 0% glycerol, 1 mM DTT and 0.03 mg/mL LMNG. The peak fractions typically m preparation of cryo-EM grids, or stored at −80°C for later use. For Drs2p ΔNC/Cdc50p, a first round of SEC had been run on a TSKg4000SW silica column on an ÄKTA purifier system at 4°C in SSR with 0% glycerol, 1 mM DTT and 0.03 mg/mL LMNG. The peak fractions were pooled and concentrated using a 50kDa cut-off centrifugal concentrator to 8 mg/mL, and was stored at −80°C for later use. A second round of SEC was run on an analytical Superdex 200 Increase 3.2/300 column on an ÄKTA purifier system at 4°C in SSR with 0% glycerol, 1 mM DTT and 0.03 mg/mL LMNG, where 50 μL sample was injected, to remove the background detergent produced by concentrating the sample. Pooling of the peak fractions resulted in a protein concentration of 0.6 mg/mL, which was used directly for preparation of cryo-EM grids. Representative chromatograms and gels are shown in Supplementary Data Figure 1A-C.

#### Activity measurement on purified protein for structural studies

The activity of the purified Drs2p-Cdc50p used for structural studies was assayed using an arsenic-based Baginski Assay^50^, a colorimetric assay for free inorganic phosphate. Drs2p-Cdc50p in LMNG to a final concentration of 10 μg/mL was added to a reaction buffer of SSR with 0% glycerol, 1mM DTT, 0.02 mg/mL LMNG and 5 mM NaN_3_ (final concentrations) as well as PS C(8:0), Brain PI4P and BeF_3_^−^(BeSO_4_ and KF in a 1:20 molar ratio) were added, when present, to final concentrations of 78 μg/mL, 20 μg/mL and 1 mM, respectively. After addition of protein to the reaction buffers, the samples were incubated on ice for 1 hour, before transfer to 30°C, and upon reaching this temperature, the reactions were initiated by addition of ATP to a concentration of 4 mM. At specific time points, 50μL sample was transferred to a 96-well microplate, and mixed with 50 μL 1:5 solution of 30mM ammonium heptamolybdate in H_2_O, and 0.17M ascorbic acid and 0.1% SDS in 0.5 M HCl. After 10 minutes at room temperature, 75 μL of arsenic solution (2% (w/v) anhydrous sodium metaarsenic, 2% (w/v) trisodium citrate dihydrate, 2% (v/v) glacial acid) was added to prevent further complexing of molybdate by phosphate. The plate was left at room temperature for 30 minutes, before measurement of the absorbance at 860 nm on a Wallac Victor 3 Multilabel plate reader (Perkin Elmer).

#### Negative Stain Electron microscopy

Copper G400-C3 grids were coated with 2% celluidine, followed by evaporation of amorphous carbon using a Leica EM SCD500 high vacuum sputter coater. Before use, the grids were glow-discharged on a PELCO easiGlow Glow Discharge Cleaning System at 25 mA for 45 seconds. 3 μL of protein sample diluted to 20 μg/mL in detergent-free buffer was applied, followed by staining three times with 3 μL 2% uranyl formate solution, which had been stored at −80°C. Micrographs were collected on a Tecnai G2 Spirit (120kV) with a Tietz F416 CCD camera using Leginon^51^. Imaging was performed at 67,000×magnification with a binned camera (pixel size 3.15 Å). Data processing including CTF-estimation, particle picking, extraction using a of 84-pixel box-size, and 2D classification was performed in cisTEM^52^ (Supplementary Data Figure 1D-F).

#### Cryo-electron microscopy

##### Sample freezing

C-flat Holey Carbon grids, CF-1.2/1.3-4C (Protochips), were glow discharged on a PELCO easiGlow Glow Discharge Cleaning System at 15 mA for 45 seconds before addition of 3μL of 0.6 mg/mL Drs2p/Cdc50p in LMNG, which had been incubated on ice for at least 1 hour with 1 mM beryllium fluoride (BeF_3_^−^), and 0.1 mg/mL Brain PI4P when indicated. The samples were vitrified on a Vitrobot IV (ThermoFisher) at 4°C and 100% humidity.

##### Data collection

The data was acquired on a Titan Krios with an X-FEG through an energy-filtered Gatan K2 camera with a calibrated pixel size of 1.077 Å/pixel at a magnification of 46,425× (MPI for Biophysics, Frankfurt). 8 second exposures fractionated into 40 frames were collected through EPU at a dose rate of 1.4 or 1.5 e^−^/Å^2^/frame, corresponding to a total dose of 56 or 60 e^−^/Å^2^.

For Drs2p ΔN104/Cdc50p 765 movies were collected on samples with 0.1 mg/mL PI4P and 3069 movies were collected without PI4P. For Drs2p ΔNC/Cdc50p 2391 movies were collected on a grid with 0.1 mg/mL PI4P and 801 movies were on a grid without additional PI4P. However, upon initial processing the datasets resulted in identical reconstructions with the same density in the PI4P-binding site (likely because of the PI4P added during the purification of this sample), and they were treated as one dataset going forward.

##### Processing

For all three datasets, movie alignment with dose weighting using all frames and contrast transfer function (CTF) determination was performed in cisTEM through Unblur^53^ and CTFFind4^54^, respectively. After manual inspection of the micrographs 2050 were selected for Drs2p ΔNC/Cdc50p, 2687 for Drs2p ΔN104/Cdc50p and 701 for Drs2p ΔN104/Cdc50p with PI4P. Using the cisTEM reference-free particle picker, a total of 1,047,615 particles were picked for Drs2p ΔNC/Cdc50p with PI4P and 2,156,578 for Drs2p ΔN104/Cdc50p and 578,440 for Drs2p ΔN104/Cdc50p with PI4P. The particles were extracted in cisTEM using a box-size of 256 pixels, and cisTEM was used for 2D classification although this was not used for selecting good particles.

##### Drs2p ΔN104/Cdc50p (E2^inhib^)

Three *ab initio* 3D references were generated in cryoSPARC^55^ from all particles, resulting in one class corresponding to the protein particle, and two corresponding to junk. Three rounds of heterogeneous 3D classification in cryoSPARC were performed, where the first round had one protein class and four junk classes, while the next rounds only used two junk classes apart from one protein class. This resulted in 769,469 particles which were subjected to heterogeneous 3D refinement, resulting in an initial reconstruction at 3.2Å from 752,881 particles.

These particles were re-extracted in Relion 3^56^ from movies aligned through the Relion implementation of MotionCor2^57^ for per-particle CTF refinement with estimation of the beam tilt and Bayesian Polishing^58^ performed in Relion 3, before the final 3D refinement resulting in an unmasked resolution of 3.0Å and a masked resolution of 2.8Å after postprocessing. Relion 3 was used for estimation of the local resolution. The processing strategy is summarized in Supplementary Data Figure 7.

##### Drs2p ΔN104/Cdc50p with PI4P (E2P^inter^)

Three *ab initio* 3D references were generated in cryoSPARC resulting in two junk and one protein-like class. These were used as reference in three rounds of heterogeneous 3D classification that resulted in 291,944 protein particles. An initial homogeneous 3D refinement of these, resulted in a reconstruction at 3.8Å of 277,569 particles. These particles were re-extracted in Relion 3 from movies aligned through MotionCor2 implemented in Relion for per-particle CTF refinement with estimation of the beam tilt and Bayesian Polishing performed in Relion 3, before the final 3D refinement resulting in an unmasked resolution of 3.4Å. However, this appeared to be a mixed state near TM10 of Drs2p, the PI4P-binding site and in parts of the autoinhibitory domain. As simple 3D classification using the refined map as reference failed to separate this heterogeneity, five new *ab initio* references were generated in cryoSPARC with similarity 1.0, to allow for very similar classes. These were then all used as references for heterogeneous 3D refinement, resulting in one class corresponding to a PI4P-bound structure, two classes identical to the determined autoinhibited structure in the absence of PI4P, and two minor junk-classes. The PI4P-bound resulted in a 3.7Å reconstruction from 78,981 particles from cryoSPARC, which was also refined in Relion 3 to an unmasked resolution of 3.9Å and a masked resolution of 3.7Å after postprocessing. Relion 3 was used for estimation of the local resolution. The processing strategy is summarized in Supplementary Data Figure 8.

##### Drs2p ΔNC/Cdc50p with PI4P (E2^active^)

Three *ab initio* 3D references were generated in cryoSPARC from all particles resulting in one class corresponding to the protein particle, and two corresponding to junk. All particles were then subjected to heterogeneous 3D classification in cryoSPARC, using each junk reference twice and the good class once, resulting in four junk classes and one with particles corresponding to the protein. This class was then subjected to heterogeneous 3D refinement in cryoSPARC resulting in an initial reconstruction at 3.4Å from 493,753 particles. The 3D refinement was repeated in Relion 2.1^59^ using a soft solvent mask, resulting in a reconstruction of similar quality. To improve the map, particle sorting was performed in Relion 2.1 and particles with a z-score above 0.9 or a defocus higher than 2.0 μm were rejected, and the remaining 418,512 were re-extracted in Relion 3 from movies aligned through Relion’s own implementation of MotionCor2. Per-particle CTF refinement with estimation of the beam tilt and Bayesian Polishing performed in Relion 3, before the final 3D refinement resulting in an unmasked resolution of 3.1Å and a masked resolution of 2.9Å after postprocessing. Relion 3 was used for estimation of the local resolution. The processing strategy is summarized in Supplementary Data Figure 9.

Data collection and processing statistics are summarized in Supplementary Data Table 1.

#### Model building and refinement

The Drs2p ΔNC/Cdc50p with PI4P was built manually in COOT^60^ guided by secondary structure predictions from RaptorX^61^, and the structures of the Na^+^/K^+^ ATPase and SERCA in the E2P state (PDB 4HYT and 3B9B) for Drs2p, which was permitted by the shared topology, and similarity of the E2P-conformations. For Cdc50p, the positions of all glycosylations were visible as at least one sugar moiety, and were used in validation of the *de novo* traced structure along with the presence of the two di-sulfide bonds. The model was refined using Namdinaor^62^ and *phenix.real_space_refine*^63^.

Drs2p ΔN104/Cdc50p was built by fitting the Drs2p ΔNC/Cdc50p model using Namdinator followed by manual editing in COOT and manual tracing of the autoregulatory C-terminus. Refinement was done using *phenix.real_space_refine*. The Drs2p ΔN104/Cdc50p with PI4P model was built based on the other two structures with manual editing in COOT followed by *phenix.real_space_refine*.

For Drs2p the 78 N-terminal residues of the construct, as well as the 46 C-terminal residues and 16 residue linker (20 in the structure of E2P^inter^) between TM10 and the autoregulatory domain in Drs2p ΔN104 were too disordered for modeling or entirely missing from the density. For Cdc50p the 19 N-terminal (18 in the structure of E2^active^) and the entire C-terminal tail of 33 residues (35 in the structure of E2^inhib^) could not be modelled.

Model validation was done through MolProbity^64^ in Phenix^65^.

Modelling and refinement statistics are summarized in Supplementary Data Table 1. Model-to-map FSC curves and representative densities from different areas of the maps are shown in Supplementary Data Figure 10.

